# The impact of channel geometry and flow regime on endothelial orientation and morphology in vessel-on-chip

**DOI:** 10.1101/2024.08.14.607791

**Authors:** Mohammad Jouybar, Sophie van der Kallen, Sheen Sahebali, Carlijn Bouten, Jaap M.J. den Toonder

## Abstract

This study investigates the impact of channel geometry and applied flow on the morphology and function of endothelial cells (ECs) within vessel-on-chip (VoC) models. Traditional organ-on-chip models often utilize rectangular cross-section channels, resulting in flat walls, sharp corners, and non-uniform wall shear stress profiles, which do not accurately mimic physiological conditions. Tubular channels with a circular cross-section provide a more in vivo-like geometry and result in a physiological uniform wall shear stress. Here, sugar 3D-printing is used for fabricating tubular channels, and stereolithography 3D-printing is applied for making rectangular channels. Using these models, the effects of both channel geometry and various flow conditions on the orientation and morphology of ECs, from both blood and lymph vessels, is explored. The research demonstrates that ECs in tubular channels exhibit a more uniform coverage and circumferential alignment/migration compared to rectangular channels. Unidirectional and bidirectional flow conditions induce alignments parallel to flow, overruling the circumferential alignment induced by curvature in tubular channels. The combination of tubular geometry and pulsatile flow induces enhanced circumferential orientation, different from rectangular channels in which cells remain primarily aligned along the flow direction under pulsatile flow. In summary, our systematic study shows that the channel curvature determines the cell orientation in static conditions, and that adding flow results in a competing effect between geometry and flow for which the resulting cell alignment depends on the flow conditions. Additionally, the study examines the initial adhesive interactions between monocytes and ECs, revealing that EC orientation induced by different flow regimes impacts monocyte rolling velocities. This finding is important for understanding immune cell motility and adhesion in healthy and diseased states. This study underlines the importance of the combined effect of channel geometry and flow conditions in VoC models, and it provides a strong motivation for the continued development of advanced OoC systems with ever more representative geometrical designs and flow control, to maximize the potential of OoC to revolutionize biomedical research and personalized medicine by providing more accurate and functional models of human physiology.

## 1 Introduction

Organ-on-Chip (OoC) models, also referred to as Microphysiological Systems (MPS), embody microfluidic-based devices featuring microchannels and compartments in which three-dimensional (3D) cell cultures are created and maintained within a microenvironment that emulates *in vivo*-like conditions within the human body. The microchannels within these models facilitate precise control over cellular and extracellular matrix (ECM) compositions, fluid transport, and mechanical cues. These unique features of OoC models make them particularly promising as disease models, offering a valuable platform for better understanding of disease development and progression, and ultimately for drug development.[1, 2]

Vessels are essential ingredients in many OoC models. The microchannels in these models are lined with endothelial cells (ECs) to represent (micro)vessels. Several technical characteristics of such a vessel-on-chip (VoC) can be controlled to mimic the physiological conditions of blood vessels, including fluid flow and channel geometry. Both these characteristics are important since they influence the behavior of blood vessels. Fluid flow induces shear stresses on the ECs cultured in microchannels, which influences the cell morphology, orientation and expression profiles.[3, 4] However, many OoC models do not represent this aspect well since they have static fluids or non-physiological flow. Moreover, the cross-sectional geometry of the perfused microchannels determines the flow profile and therefore the specific effect of the shear stress. Most OoC models have channels with rectangular cross-section channels, resulting in non-uniform shear stress profiles that are not physiological. Also, the possible effect of curvature on EC morphology and behavior is not well modeled in conventional rectangular channels. Microchannels with lumens with a circular cross-section may better mimic the physiology and cellular behavior of tubular structures like (micro)vessels and epithelial ducts. Tubular microchannels provide a more *in vivo*-like geometry and enable to generate a uniform wall shear stress under physiological flow conditions.[5, 6]

Recent studies have focused on improving the geometrical features in microfluidic devices for enhancing the physiological relevance. Several techniques have been introduced to integrate channels with a circular cross-section into OoC systems to generate tissues that contain tubular structures,[7] such as (micro)vessels.[8, 9] Early techniques include photoresist reflow, viscous finger patterning,[10, 11] and template-casting.[12, 13, 14, 15, 9] Li et al. fabricated a network of cylindrical channels using a resist reflow technique and seeded ECs in these channels to generate blood vessels.[16] de Graaf et al. utilized the viscous finger patterning technique to generate a tubular channel with an average diameter of ∼ 330 µm within a collagen I scaffold. When hiPSC-ECs were seeded in these channels, they formed microvessels.[11] Various types of templates have been used in the template casting method. Jiménez-Torres et al. used polydimethylsiloxane (PDMS) rods to create individual tubular channels and bifurcations, which were lined with endothelial cells.[14, 13] In other studies, micro-needles were used to generate two tubular channels in close vicinity. For example, Nguyen et al. generated parallel channels this way, one lined with ECs and the other containing cancer cells.[15] In another study, Kutys et al. developed a vascular and breast duct model by combining two tubular channels, one forming a perfused channel and the other forming a dead-ended channel positioned perpendicularly 500 µm away from the first channel.[9] For realizing more intricate tubular geometries, sacrificial materials were 3D-printed as templates for obtaining tubular channels embedded in hydrogel or in cured polymer.[17, 18, 19, 20, 21] For example, carbohydrate glass fibers were printed as sacrificial molds to generate perfusable patterned tubular networks.[18, 20] Miller et al. utilized this technique to obtain 3D vascular networks in various types of hydrogels.[18] Recently, Nie et al. introduced a method to create tubular OoC models combining a novel cell seeding technique with 3D-printing of sacrificial fiber; they demonstrated the method with a perfusable renal proximal tubule-on-a-chip with physiological dimensions.[22] In some other studies, bioprinting inks (such as pluronic F-127 and gelatin) were used for 3D printing the sacrificial mold.[19, 21] For example, Kolesky et al. fabricated a thick vascularized tissue including a network of endothelialized tubular channels enabling long-term perfusion.[19] In a recent study, Zhang et al. fabricated a system of two parallel lumens using 3D printing of gelatin as sacrificial mold;[21] this system incorporated duo-recirculating flow for culturing vascularized renal proximal tubules with glucose re-absorption function.

In addition, laser ablation techniques (such as nano- and femto-second laser machining) have been recently utilized for the integration of tubular channels in OoC models.[23, 24, 25] In a recent study, Enrico et al. applied femtosecond laser machining to obtain tubular channels lined with ECs in close vicinity to glioblastoma spheroids.[25] While these methods have significantly advanced the capacity to control geometric cues in OoC systems, the application of flow conditions was not always implemented in these studies.

Several studies have looked into the effect of flow on endothelium morphology in various types of 2D and 3D models (Supplementary table 1). Many of these previous studies showed that ECs aligned in the direction of the flow under unidirectional flow.[3, 4] In contrast, some studies reported that ECs did not organize to align in the direction of the flow when exposed to irregular or pulsatile flow. [26, 27, 28] Gravity-driven microfluidic devices were placed on slow-tilting tables to generate uni- and bi-directional flow.[21, 29]

Yang et al. showed that ECs aligned in the direction of unidirectional flow, and that alignment drastically reduced in bidirectional flow. Some recent studies applied unidirectional and bidirectional flow in tubular channels. Zhang et al. showed that the majority of HUVECs were aligned at an angle ∼ 30°-60° with respect to the flow direction in both uni- and bi-directional flow conditions.[21] In another study, Gong et al. showed that most of the cells were oriented in the flow direction, however, many cells still aligned in the ∼ 30°-60° or ∼ 100°-120° direction under non-continuous dynamic flow.[30] While different types of flow conditions in OoC systems were shown to affect the morphology and behavior of ECs, a systematic study of the influence of different flow conditions on the morphology and orientation of both blood and/or lymphatic ECs is lacking. In addition, a systematic study of the combined effect of geometry (tubular versus rectangular) and flow on endothelium morphology and function remains unavailable.

The morphology of ECs plays a crucial role in their interactions with immune cells.[31] Monocytes are a subgroup of leukocytes which have the ability to transmigrate from the blood stream into surrounding tissues and differentiate into macrophages or dendritic cells as part of the inflammatory response against foreign bodies and during tissue damage.[32] ECs respond to blood flow by sensing the fluid flow-induced shear stress on the vascular wall[33] and a variety of responsive proteins, such as integrins and adhesion molecules, activate multiple signaling networks.[34] The adhesive interactions between monocytes and ECs, via dynamic receptor-ligand bonds, facilitate monocyte rolling behavior preceding monocyte arrest and adhesion to the endothelium. Many factors play role in this interactive phenomenon. Even though sub-cellular components such as endothelial integrins (e.g. via association with VCAM-1 and ICAM-1) and selectins have been the main focus of studies, cell morphology, orientation, and deformability have been suggested to influence this adhesive behavior.[35, 36, 37, 31] Some studies indicate that physiochemical properties such as bond interaction rate, dictated by the flow, primarily determine the adhesion behavior.[38, 39] Further investigations are needed to unveil the mechanistic aspects of the interactions between ECs and immune cells.

Here, we conducted a systematic study of the effects of channel geometry and flow conditions in microfluidic channels in VoC models. We investigated the morphology and orientation of blood and lymphatic Ecs lining the walls of tubular channels versus conventional channels with a rectangular cross-section. Moreover, the combined effect of channel geometry and flow was studied by applying various flow conditions, including static, bidirectional, unidirectional, and pulsatile flow in circular and rectangular cross-section channels.

We utilized sugar glass 3D-printing to fabricate tubular microchannels, and stereolithography(SLA) 3D-printing to fabricate rectangular microchannels. To investigate the influence of cellular orientation, resulting from different flow conditions, on immune cell motility and adhesion, we studied the initial adhesive behavior of circulating monocytes on different endothelial alignments by analyzing monocyte rolling velocities on the endothelium. This study enhances our basic understanding of the importance of channel geometry and hydrodynamic conditions on cellular orientation and morphology in VoC models. Finally, our results show how the orientation of ECs can affect monocyte motility in healthy versus disease conditions.

## 2 Results and Discussion

### 2.1 Fabrication of circular and rectangular cross-section channels

We used the sugar 3D-printing technique previously described[20, 22] to obtain channels with a circular cross-section (Figure 1 a-c). Sugar glass structures were printed using a dedicated extrusion-based 3D printer, forming a sacrificial mold, and then the structures were cast in PDMS (Figure 1 a). After the PDMS was cross-linked and the sugar glass was selectively dissolved, the sugar fibers were removed by dissolving in water overnight to form hollow tubular channels (Figure 1 b). The PDMS slab containing the tubular lumen was assembled with a top layer with reservoirs aligned to the inlet and outlet of the channel (Figure 1 c). The reservoirs were used for media exchange, designed to limit evaporation from the channel. The channels with a rectangular cross-section were fabricated by casting PDMS on a SLA 3D-printed mold and subsequent curing; the PDMS slab containing the channel was sealed with a top layer including reservoirs (Figure 1 d). We fabricated tubular channels with a circular cross-section with a 300 µm diameter, and rectangular channels with a cross-section of 188 µm x 376 µm (*h* × *w*), to have the same cross-sectional area; the channel also have equal lengths. Endothelial cells were cultured in these channels for 3 days in static conditions to cover the walls of the channels (Figure 1 e). Three different flow conditions were applied for further 3 days and then samples were analysed for cellular morphology and alignment (Figure 1 f).

**Figure 1:**
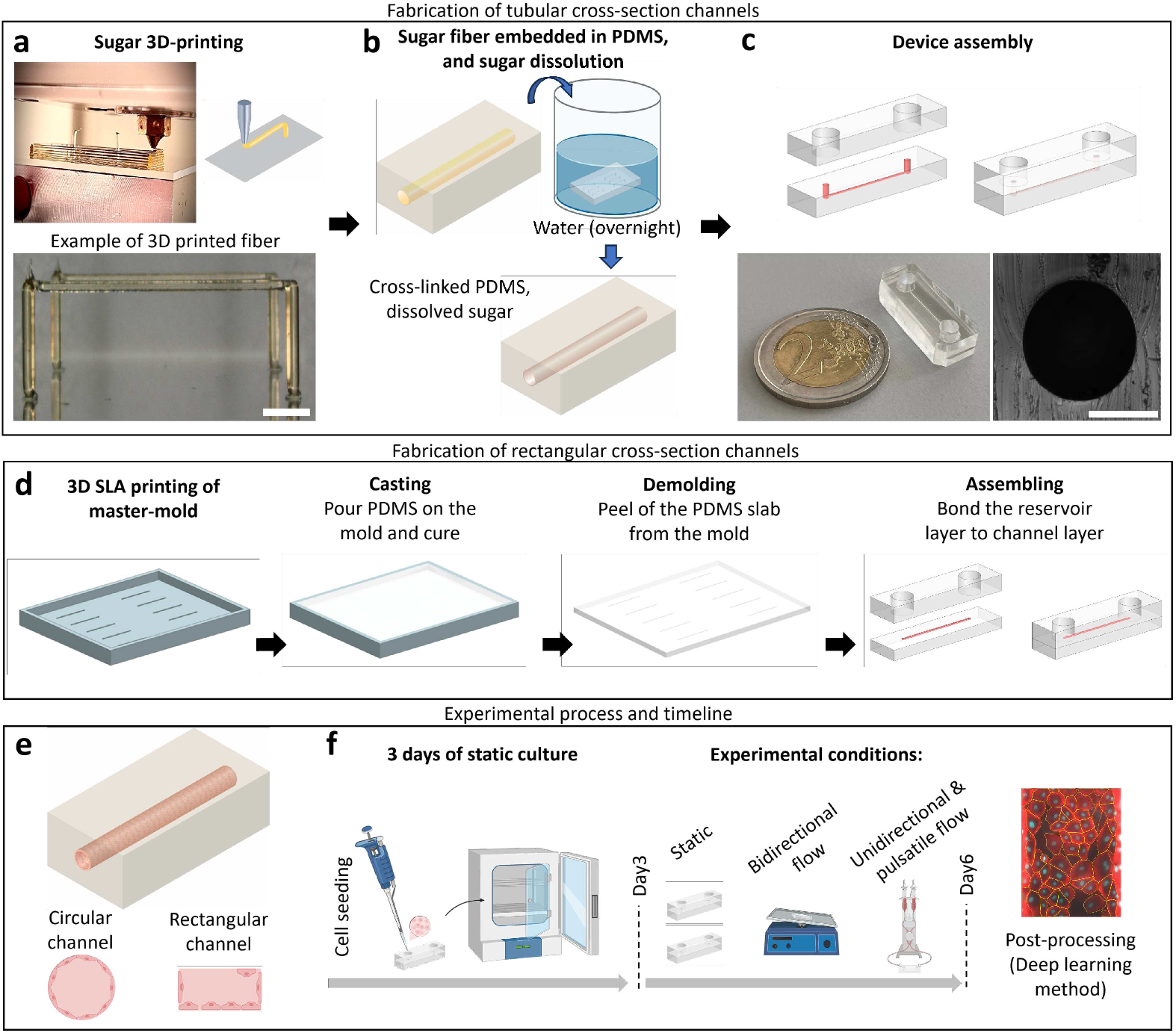
Fabrication methods and experimental process. a-c) Fabrication process of tubular channels. (a) Image and schematic illustration of a dedicated extrusion-based 3D-printer while printing sugar fibers, and images of printed sugar fibers suspended between pillars; scale bar: 1 mm. b) PDMS casting around the sugar fiber and sugar dissolution after PDMS cross-linking. c) Schematic illustration of PDMS device assembly, and images of the PDMS chip and the channel cross-section; scale bar: 200 µm. d) Fabrication process of channels with a rectangular cross-section, consisiting of stereolithography (SLA) 3D-printing of molds, casting in PDMS and curing, demolding the PDMS layer, and assembly with a top layer including in- and outlet reservoirs. e) Schematic representation of cell seeding in tubular and rectangular channels for generating vessel-on-chip systems. f) Schematic representation of the experimental process; samples were kept in culture in static (no flow) conditions for 3 days, and then different flow conditions (bidirectional, unidirectional, and pulsatile) were applied for 3 days. Samples then were fixed, imaged and processed.

In the current work, the cross-sectional dimensions of our channels are a few hundred micrometers, which is in the range of common sizes of microfluidic channels, and we do not investigate the effect of channel dimensions. This is a relevant topic for future work; for example, Kim et al. showed in a numerical study that the diameter of capillary lumens influences the migration direction of the cells.[40] With our sugar 3D-printing technique, it is possible to realize smaller channels, down to 50 µm, by using smaller extrusion nozzle tips.[20, 22] The SLA 3D-printing technique we used to make rectangular channels is limited in resolution (25 µm); hence, other techniques such as photo-lithography would be beneficial for fabricating rectangular channels with smaller dimensions.

Another factor that could influence the results of the current study is the material stiffness of the channel. The microfluidic channels in OoC systems are mostly made either in polymers (such as PDMS, with a Young’s modulus of around 1 MPa) or in hydrogels (with a Young’s modulus typically around 1 kPa). It has been shown that the substrate stiffness influences cell attachment, migration, and morphology.[41, 42, 43, 44, 45, 46] In this study, we used PDMS devices while controlling cell adhesion by pre-treatment of the channel surfaces with fibronectin prior to cell seeding. This thin coated layer works as the interface between cells and channel walls, which possibly mediates the cell migration.[40] While focusing on the effect of geometry and flow in the current study, our techniques can be utilized in future work to study the effect of different materials and associated stiffness on the cell behavior.[22]

### 2.2 Tubular geometry of channels induces a circumferential alignment and migration of endothelial cells

The distribution and morphology of HUVECs in microchannels with circular and rectangular cross-sections was initially studied after 6 days of static culture with daily media refreshment. We noticed that ECs uniformly covered the whole surface of the tubular channel. On the other hand, the top surface of the rectangular cross-section channel was not fully covered by cells (Figure 2 a-d); the number of cells residing on the bottom surface of the rectangular channel was substantially higher than at the top (Figure 2 e). We observed this phenomenon in other culture conditions as well, including bidirectional, unidirectional and pulsatile flow (Figure 2 e). In this study, we seeded HUVECs with the chip in upright orientation, and we subsequentially rotated the chip 90° to both sides; hence the difference in surface distribution emerges from a difference in migration caused by channel geometry. With several previously reported devices including rectangular cross-section channels, it was necessary to flip the chip 180° during cell seeding to achieve homogeneous cell attachment to all walls of the channel.[47, 48] We intentionally avoided 180° rotation to observe if cells could naturally reach the top of the channel by migration, resulting in full coverage only for the tubular channel. The corners of the rectangular channels most probably hindered cell migration from the bottom surface to the side walls and the top surface. Even if all walls of rectangular channels are covered by cells using the 180° flipping method, the presence of the corners will probably influence the cellular behavior, morphology, and migration pattern during the culture period, whereas the curved surfaces of the tubular channels facilitate the migration and interaction of the cells over the entire cross-section under all conditions.[45, 40, 29, 30]

**Figure 2:**
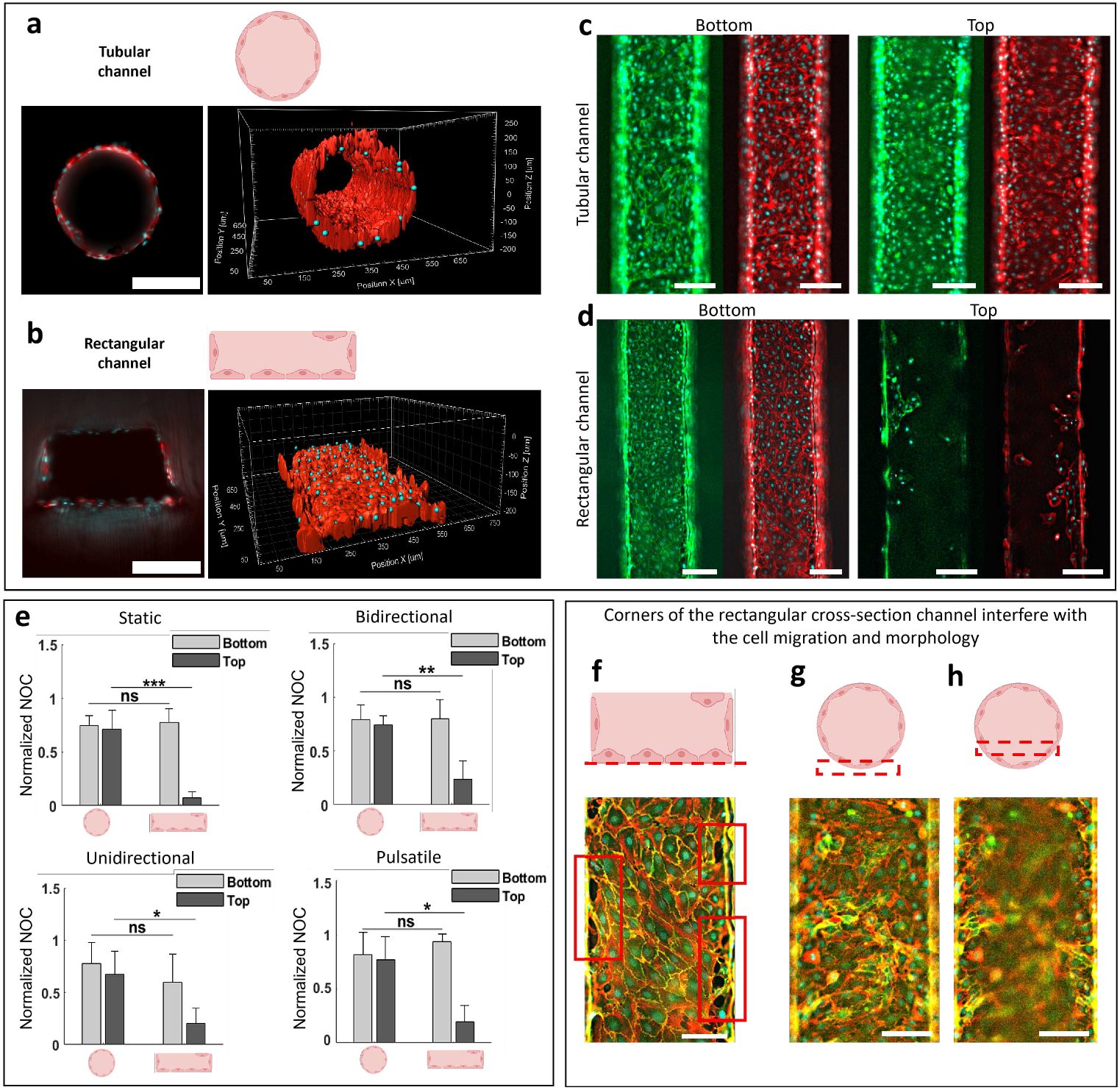
Comparison of the distribution of ECs in channels with circular and rectangular cross-sections under different flow conditions. a, b) Schematic and image of cross-section and rendered 3D construction of ECs lining the a) tubular and b) rectangular channels. Cell nuclei are shown in cyan, and F-actin is shown in red. In the rendered images the red is reconstructed from the F-actin signal, and the cyan dots from the cell nuclei signal.Scale bar, 100 µm. c-d) HUVECs stained for cell nuclei (cyan), CD31 (green) and F-actin (red), after six days of static culture, in the bottom and top of the (c) tubular and (d) rectangular channels. Scale bar, 200 µm. e) Average normalized cell count in the bottom and top regions of the tubular versus rectangular channels cultured under static, bidirectional, unidirectional and pulsatile flow conditions. Normalized values of number of cells in each culture condition (Normalized NOC) were calculated separately for each condition by dividing the values by the highest number of cells observed in the particular culture condition. f-h) The HUVEC layer at the (f) bottom of the rectangular channel, and at the g, h) bottom of the tubular channel. The figures include the combined fluorescent signals of cell nuclei (cyan), CD31 (green), and F-actin (red). The red rectangles in f) show the empty gaps at the edges of the rectangular cross-section channels. Scale bar, 100 µm. *P <0.05, **P <0.01, and ***P <0.001 indicate statistical significance. Paired t-tests were done to determine statistical significance.

Fluorescent images of the HUVECs sheet, stained for F-actin and CD31, revealed distinct empty regions along the edges of the bottom of the rectangular channels (Figure 2 f and Supplementary Figure 1); such gaps were not observed in the images of cells in the bottom and middle of the tubular channel (Figure 2 g-h). Moreover, F-actin stress fibers appeared more intensely in the cells around the gaps (Supplementary Figure 1), showing that the prominence of stress fibers in cells was influenced by the geometrical cues from the corners of rectangular channels. In addition, cell tracking analysis of live time-lapse images at the bottom of the channels revealed a consistent circumferential movement of cells along the inner surface of the tubular channel throughout the cell culture period (Supplementary Video 1 and Supplementary Figure 2 a). In contrast, the motion of the cells on the bottom wall of the rectangular channel lacked a consistent direction (Supplementary Figure 2 b). The average migration angle of the cells, i.e. the angle of the line connecting the starting and final locations of the cells over a period of 20 hours with respect to the line parallel to the channel main axis, was close to 80° in tubular channels, in line with the observed circumferential motion; in rectangular channels, the average migration angle was 45° with a very wide spread (Supplementary Figure 2 c). Finally, cells moved along a straighter path in tubular channels as compared with rectangular channels, quantified by the ratio of migration distance over total path length in Supplementary Figure 2 d.

Previous studies confirm that channel geometry strongly affects the cellular morphology, behavior, motility, and barrier properties of endothelial layers in Vessel-on-Chip models.[45, 49, 50, 51, 46] These findings are in line with the growing body of evidence that confirms that surface curvature influences the spatiotemporal organization of cells and tissues cultured on surfaces.[52, 53, 40] These geometry-induced effects can have important implications - for example, vessel curvature can affect the intra-endothelial tension; when the cells need to adapt to curvature, the cytoskeleton and junctions are under tension, possibly facilitating intravasation and cellular traverse.[45, 54] In agreement with earlier works, our results underline that the use of channels with circular cross-sections is essential for obtaining a good representation of tubular biological structures such as blood vessels, irrespective of the influence of flow which we will discuss next.

### 2.3 Effect of flow conditions on the alignment and morphology of HUVECs in circular and rectangular cross-section channels

Several recent studies have investigated the effect of flow on endothelial cell alignment, including *in vivo* and *in vitro* testing, and using various approaches including channels with circular and rectangular cross-sections (Supplementary Table 1). However, the combined effects of flow and channel geometry on the morphology of ECs has still not been studied and reported consistently. We present a systematic study of the influence of different flow conditions on EC morphology for both circular and rectangular cross-section channels. After seeding the cells in the channels, they were left to grow in static conditions for three days, after which different flow conditions were applied for an additional 3 days, including static, bidirectional, unidirectional, and pulsatile flow. These conditions were chosen because the unidirectonal and pulsatile flow are typical conditions found in vascular system.[55] The bidirectional flow condition was added because this is normally applied by commercial rockers to induce perfusion in OoC systems, which have become fairly common recently. Then, immunofluorescent images were made for morphological studies (Figure 2 f). The results are shown in Figures 3 and 4. Under static culture conditions, the cells aligned mostly perpendicularly to the main axis of channel in tubular channels, i.e., they exhibited circumferential alignment, while the majority of cells within the rectangular channels aligned along the channel’s main axis (Figure 3 a and Figure 4 a), probably because of the presence of the edges. Yet, many cells aligned at 30-150° with respect to the main axis of the channel. In static culture, the only shear stress was applied by daily media refreshment. The maximum wall shear stress resulting from hydrostatic pressure during media refreshment, was estimated as *τ*_max_ ≃ 0.25 Pa in the tubular channel and *τ*_max_ ≃ 0.28 Pa in the rectangular channel. In some studies, carrying out regular several media refreshments was considered as a dynamic culture,[30] however, we call this a static culture condition because the values of the shear stress are low and it occurs just once a day. Note, that the wall shear stress in tubular channels is spatially uniform, whereas it varies in space for rectangular channels; the wall shear stress values represent averages in space.

**Figure 3:**
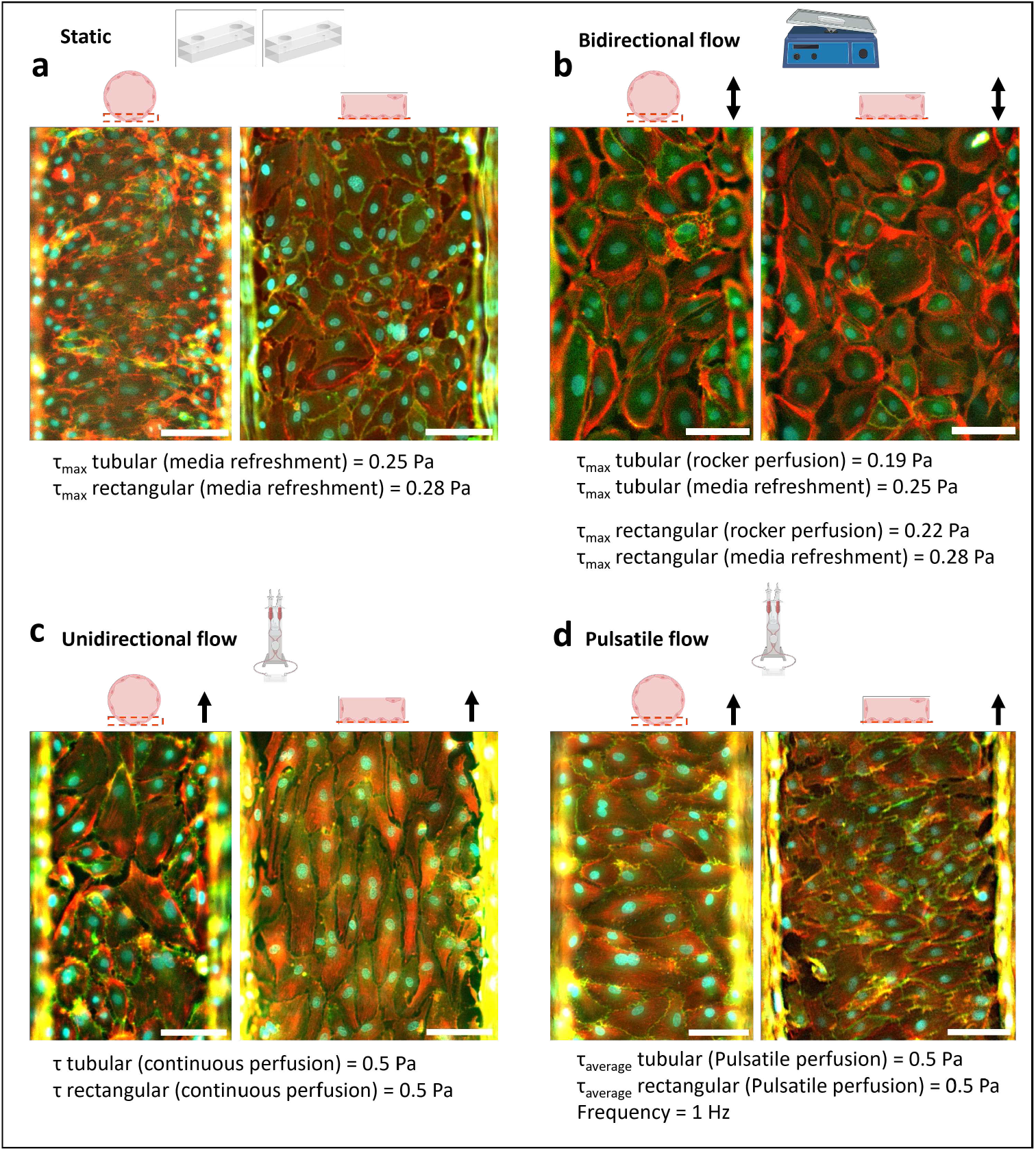
Endothelial cell alignment and morphology under different flow conditions in tubular and conventional rectangular channels. a-d) Images of HUVECs at the bottom of the tubular and rectangular channels in a) static conditions (no flow), b) bidirectional, c) unidirectional, and d) pulsatile flow conditions. Images show overlay stainings of F-actin in red, CD31 in green, and cell nuclei in cyan; arrows show the flow direction along the main axis of the channels. *τ*_*max*_ is the maximum wall shear stress applied on the ECs lining the walls of the channels (circular and rectangular cross-section) resulting from daily media refreshment or from cyclic motion of the rocker; the wall shear stress value was constant in unidirectional flow (*τ*), and it was cyclic in pulsatile flow (*τ*_*average*_ with an on-off frequency of 1 Hz). Note, that the wall shear stress in tubular channels is spatially uniform, whereas it varies in space for rectangular channels; the wall shear stress values represent averages in space. scale bar: 100 µm.

**Figure 4:**
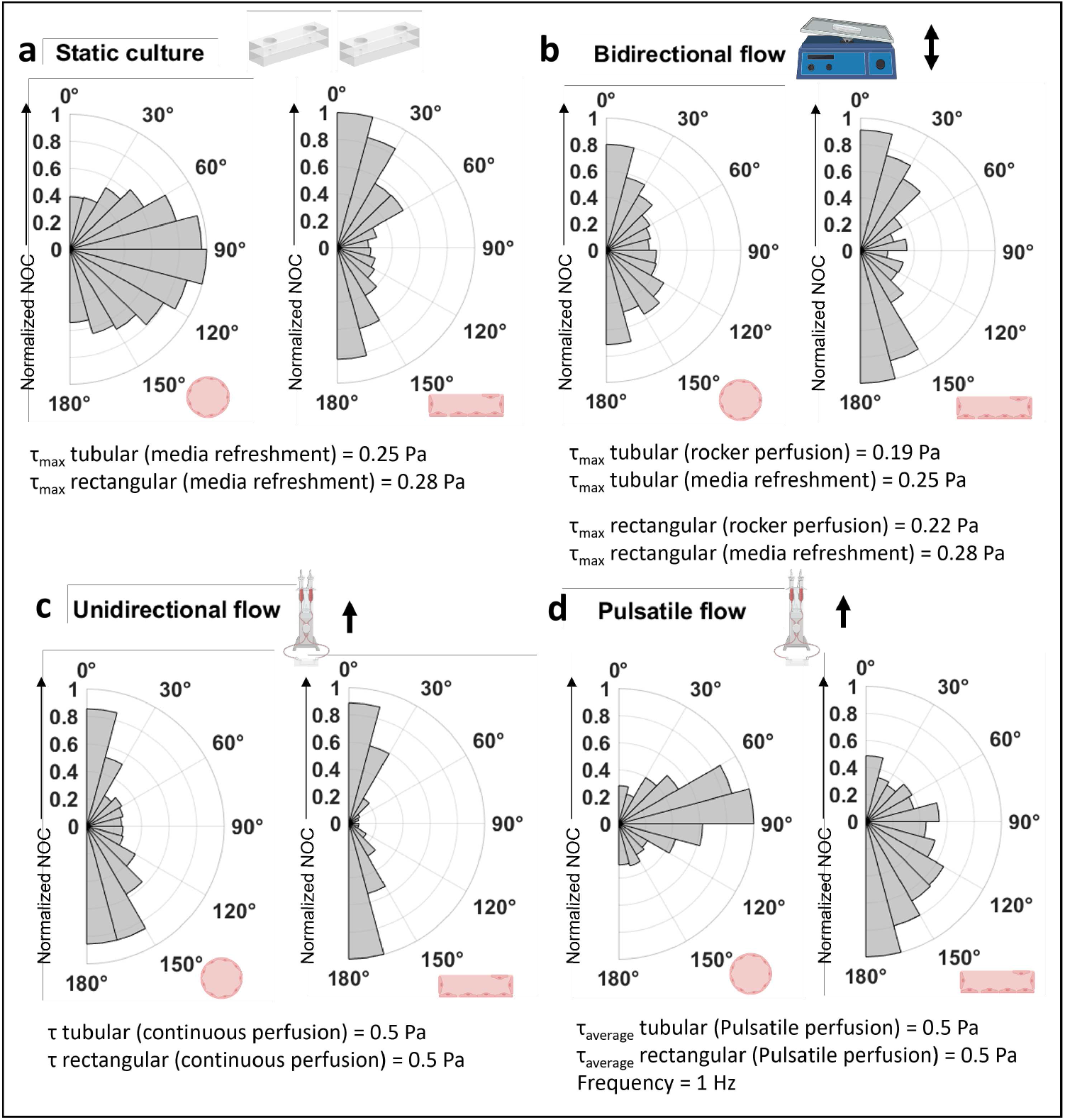
The orientation of HUVECs under different flow conditions in tubular and rectangular channels. a-d) Polar histograms representing the orientation of cells in channels with tubular versus rectangular cross-sections in a) static, b) bidirectional, c) unidirectional, and(d) pulsatile flow conditions; the radius of the slice bars represents the average of normalized number of cells (Normalized NOC). To normalize the data, the number of cells in each condition was divided by the highest number of cells observed in the culture condition. This normalization process allows for a comparative analysis of cell distribution across different conditions, ensuring that the values are on a consistent scale; 90° alignment corresponds to perpendicular orientation to the flow, while 0° and 180° alignment correspond to orientation along the main axes of channels, shown by the arrows beside the radius of polar histograms. The small arrows and schematics represent the flow conditions. *τ*_*max*_ is the maximum wall shear stress applied on the ECs lining the walls of the channels (circular and rectangular cross-section) resulting from daily media refreshment or cyclic motion of the rocker; the wall shear stress value was constant in unidirectional flow (*τ*), and it was cyclic in pulsatile flow (*τ*_*average*_ with an on-off frequency of 1 Hz). Note, that the wall shear stress in tubular channels is spatially uniform, whereas it varies in space for rectangular channels; the wall shear stress values represent averages in space.

In bidirectional and unidirectional flow conditions (Figure 3 b, c and Figure 4 b, c), the cells primarily aligned in the main channel direction, i.e., in the direction of the flow, for both channel geometries. Clearly, in these cases, the circumferential alignment induced by the tubular channel geometry seen in the static culture, is overruled by the effect of flow; this effect is most prominent for unidirectional flow. The strongest alignment with the main flow direction was observed for the cells under unidirectional flow in the rectangular channel for which geometrical and flow induced cues go hand in hand. In the bidirectional flow condition, the cyclic wall shear stress was applied using a rocker tilting to a maximum angle of ±15°at 5 cycles per minute. The maximum wall shear stress in the rocking cycles was *τ*_max_ ≃ 0.19 Pa in the tubular channel, and *τ*_max_ ≃ 0.24 Pa in the rectangular channel, i.e., lower than the typical physiological wall shear stress of 0.5 Pa. Moreover, the shear stress resulted from daily media refreshment, like the static culture, was also applied in the bidirectional flow condition. In the unidirectional flow condition, a wall shear stress of *τ*=0.5 Pa on the channel walls, comparable with that in physiological conditions, was applied by setting the flow rate.[55]

Remarkably, in pulsatile flow, the ECs were aligned in the circumferential direction in tubular channels, whereas the alignment in the rectangular channels was more ambiguous, with a significant proportion of cells aligning in the main channel direction (Figure 3 d and (Figure 4 d). The latter can be explained by the aligning effect of the channel edges. Like for the unidirectional flow, an average wall shear stress of *τ*_average_ = 0.5 Pa was applied in the pulsatile flow condition; the switching frequency was set at 1 Hz to generate a pulsatile flow altering periodically between flow and no flow conditions.

The images in Figure 3 show that the expression of actin was influenced by the flow conditions. In static and bidirectional flow conditions, actin was expressed intenser at the cell borders, however, in unidirectional and pulsatile flow conditions, actin fibers were more expressed throughout the cells.

Our results are in line with previous studies that showed that ECs align in the flow direction under unidirectional flow.[3, 4] Some studies reported that ECs did not align with the flow direction when exposed to irregular or pulsatile flow, to which our results for pulsatile flow partly agree.[26, 27, 28] Several studies used gravity-driven systems to generate unidirectional and bidirectional flow through microfluidic channels with rectangular and circular cross-section.[21, 29] Of these studies, Yang et al. showed that ECs within a rectangular cross-section channel oriented in the direction of unidirectional flow, and that this alignment reduced under bidirectional flow.[29] The applied wall shear stress was comparable in our work; the reduced alignment in the bidirectional flow condition is due to the cyclic oscillation of shear stress value. In another study with a tubular channel geometry, Zhang et al. showed that the majority of HUVECs were aligned at ∼ 30°-60° with respect to the flow direction under gravity-driven unidirectional and bidirectional flow.[21] This result is similar to ours for bidirectional flow in a tubular channel (Figure 4 b-left), with a comparable applied wall shear stress (0.1-0.2 Pa); the wall shear stress applied in our unidirectional flow was higher than that applied by Zhang et al., which explains the stronger alignment along the flow. Along with these studies, our results show that EC alignment results from the combination of flow conditions and channel geometry. In particular for pulsatile flow, the difference in cell alignment between channels with circular and rectangular cross-sections is notable. The curvature of the tubular channels led to a decrease of the number of flow aligned cells under unidirectional and bidirectional flow compared to rectangular channels, and it promoted the circumferential alignment in pulsatile flow, whereas the alignment in the rectangular channel was more ambiguous for this condition. In conclusion, our systematic study by including both the channel geometry and flow conditions showed that the channel curvature dominated the cell orientation in static condition, and affected the cell alignment under flow conditions.

In addition to the alignment, we analyzed the morphological characteristics of the ECs, in particular the aspect ratio, circularity, and area of single cells (Figure 5). The aspect ratio of HUVECs covering the bottom of the channels was the highest for pulsatile flow in the tubular channel, and for unidirectional flow in rectangular channel (Figure 5 a). Interestingly, bidirectional flow did not alter the aspect ratio of cells with respect to the static condition (Figure 5 a). Unidirectional flow increased the aspect ratio slightly in tubular and drastically in the rectangular channel with respect to bidirectional flow. On the other hand, pulsatile flow increased only the aspect ratio in tubular channels (Figure 5 a). Bidirectional flow induced the highest circularity in HUVECs in both tubular and rectangular channels (Figure 5 b). The circularity was lower in unidirectional and pulsatile flow in comparison with bidirectional flow for both tubular and rectangular channels; the circularity was lowest under pulsatile flow in the tubular channel and under unidirectional flow in the rectangular channel (Figure 5 b). Clearly, our results show that the same flow conditions (in particular for unidirectional and pulsatile flow) induced different cell morphology in tubular than in rectangular channels, highlighting the importance of channel geometry on cellular morphology. The normalized single cell area increased in all flow conditions in comparison with the static culture condition, except for unidirectional flow in the tubular channel (Figure 5 c). The area of cells was remarkably higher under pulsatile flow than for other flow conditions in both types of channels. Although there are some studies of the orientation of the cells under different flow conditions,[56, 57, 33, 26, 27, 28] a systematic study of the morphological features as presented here has not been reported before. Recent studies mostly reported the increased aspect ratio of cells under pulsating or continuous flow conditions, by measuring the shape index.[56, 57]

**Figure 5:**
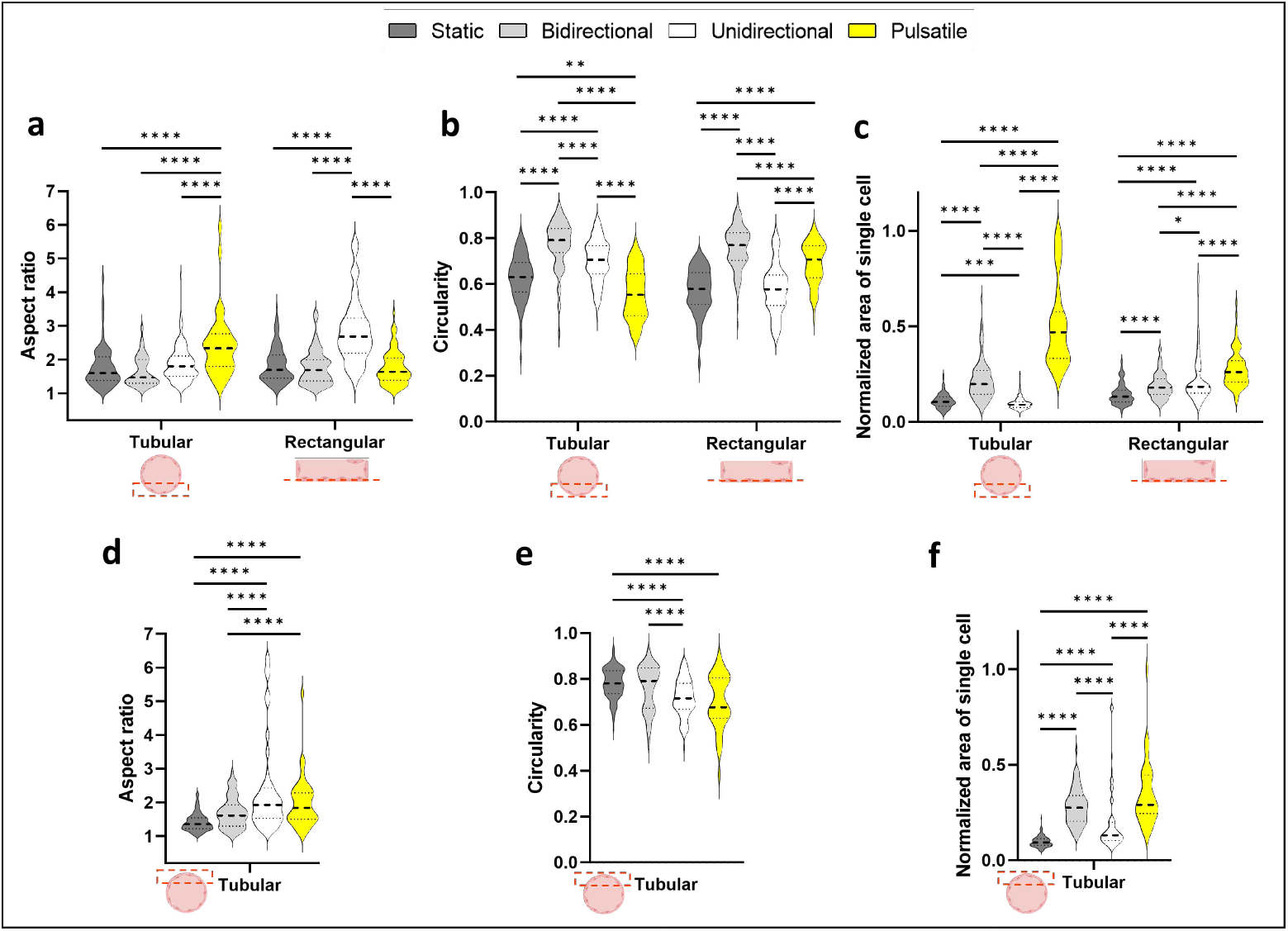
Morphological characteristics of HUVECs for endothelium cultured in tubular and rectangular channels under different flow conditions. a-c) Violin plots, representing the (a) aspect ratio, (b) circularity, and (c) normalized area of single cells at the bottom of channels with circular and rectangular cross-section channels under four different flow conditions, including static, bidirectional, unidirectional, and pulsatile flow. d-f) Violin plots, representing the (d) aspect ratio, (e) circularity, and (f) normalized area of single cells near the top of tubular channels with circular cross-section channels under four different flow conditions, including static, bidirectional, unidirectional, and pulsatile flow. To normalize the data for single cell area, the area of the cells was divided by the highest area value among all culture conditions. *P <0.05, ***P <0.001, ****P <0.0001 indicate statistical significance. ns indicates non-significance. One-way ANOVA, followed by Tukey post-hoc analysis were done to determine statistical significance.

The morphological analysis of cells residing in the top region of the tubular channel gave results similar to those of the bottom surface, with a few exceptions (Figure 5 d-f). First, the aspect ratio was highest for unidirectional flow (Figure 5 d), rather than for pulsatile flow as at the bottom of the channel (Figure 5 a). Nevertheless, it appeared that unidirectional and pulsatile flow conditions increased the aspect ratio and diminished circularity both at the bottom and the top of the tubular channel (Figure 5 a, b, d, e). Secondly, while the bidirectional and pulsatile flow conditions induced the largest cell areas on both the bottom and top surfaces, for pulsatile flow the cell area was smaller on the top surface compared to the bottom (Figure 5 c, f). Our data suggest a direct relation between the increased single cell area with the cyclic flow conditions (bidirectional and pulsatile) (Figure 5 c, f). Previous studies reported that unsteady flow conditions influence cell shape and morphology through altered actin cytoskeleton organization.[58, 59, 60] Nevertheless, we observed a different actin expression in the bidirectional versus the pulsatile flow conditions; actin was more expressed at the borders of the cells in bidirectional flow, but it spread throughout the cell body in pulsatile flow (Figure 3 b, d). This might be the result of the different wall shear stress values in these cases, and not of the cyclic flow regime.

### 2.4 Effect of tubular versus rectangular channel geometry on lymphatic endothelial cells under static and flow culture conditions

We conducted experiments using an alternative cell type to HUVECs, namely human dermal lymphatic endothelial cells (HDLECs), and we analyzed the effect of channel geometry and flow conditions on cell alignment and morphology (Figure 6). While direct comparative studies of cell behavior in tubular versus rectangular channels for lymphatic vessels are limited, existing research suggests that lymphatic endothelial cells (LECs) and blood ECs (here, we use HUVECs) exhibit distinct motility behaviors due to their different functional roles and molecular characteristics.[61, 62, 63] Moreover, while extensive research has been done on blood ECs in OoC systems, lymphatic vessels are often neglected in organ microenvironment and disease modeling systems. Recent studies began to include lymphatic vessels in OoC systems considering their relevance in the progression of various diseases, including metastasis.[2, 64, 30, 65] These arguments support our choice to analyse the organization of lymphatic ECsin our VoC systems. Given that the lymphatic circulatory system lacks the high flow rates and related wall shear stresses typical for the blood circulation,[66, 55] we only investigated static and bidirectional flow conditions. Recently, various types of rocker pumps have become commercially available to realize bidirectional flow conditions. These systems are commonly utilized for passive flow perfusion in OoC applications, often without considering their impact on cellular morphology.

**Figure 6:**
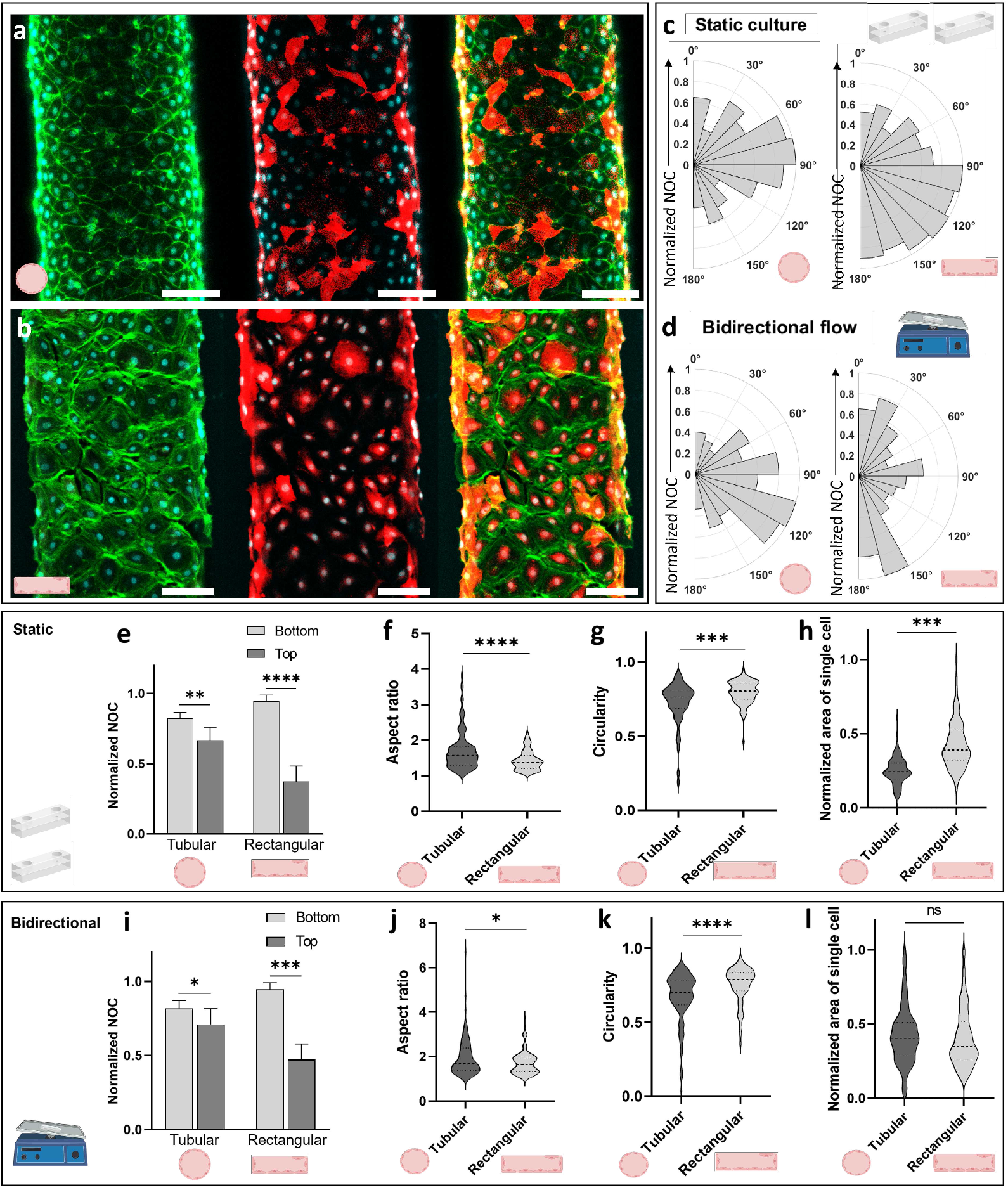
Effect of flow conditions and channel geometry on the alignment and morphology of lymphatic endothelial cells. a, b) Images of HDLECs at the bottom of the (a) tubular and (b) rectangular channel. From left to right, images show staining of F-actin in green, LYVE1 in red, and overlap of both. Cell nuclei are stained in cyan; scale bar: 100 µm. c, d) Polar histograms representing the orientation of cells in channels with tubular versus rectangular cross-sections in (c) static and (d) bidirectional flow conditions; the radius of the slice bars represents the average of normalized number of cells (Normalized NOC). To normalize the data, the number of cells in each condition was divided by the highest number of cells observed in the culture condition. This normalization process allows for a comparative analysis of cell distribution across different conditions, ensuring that the values are on a consistent scale. 90° alignment corresponds to orientation perpendicular to the flow direction, while 0° and 180° alignment correspond to orientation along the main axes of the channels, shown by the arrows beside the radius of polar histograms. e, i) Average of normalized number of cells (divided by the highest number of cells in each condition) at the bottom and top of the tubular and rectangular channels under (e) static and (i) bidirectional flow conditions. f-h, j-l) Violin plots, representing the morphological characteristics in tubular and rectangular channels, in terms of (f, j) aspect ratio, (g, k) circularity, and (h, l) normalized area of single cells under (f-h) static and (j-l) bidirectional flow conditions. *P <0.05, **P <0.01, ***P <0.001, ****P <0.0001 indicate statistical significance. ns indicates non-significance. Paired t-tests were done to determine statistical significance.

The same low values of wall shear stresses as for HUVECs were applied for HDLECs. In static culture, the maximum wall shear stress, resulting from hydrostatic pressure during media refreshment, was *τ*_max_ ≃ 0.25 Pa in the tubular channel and *τ*_max_ ≃ 0.31 Pa in the rectangular channel. In some studies, applying several media refreshments was considered as a dynamic culture,[30] however, we call this a static culture condition because the values of the wall shear stress are low and it occurs just once a day. In bidirectional flow, the maximum wall shear stress on the channel walls in the rocking cycles was *τ*_max_ ≃ 0.19 Pa in the tubular channel, and *τ*_max_ ≃ 0.24 Pa in the rectangular channel (tilting to a maximum angle of ± 15°at 5 cycles per minute. Moreover, the wall shear stress resulting from daily media refreshment, like in the static culture, was also applied in the bidirectional flow condition.

Similar to HUVECs, we observed distinct morphological characteristics of HDLECs grown in circular and rectangular channels. Actin expression of HDLECs was not very different in tubular versus rectangular cross-section channels, (Figure 6 a,b) as we observed the same for HUVECs (Figure 3 a, b). In addition to actin, HDLECs were also stained for LYVE-1, a marker for lymphatic endothelial cells. We noticed variations and inhomogeneities in LYVE-1 expression (Figure 6 a,b); these endothelial cells were separated from blood endothelial cells based on the positive marker podoplanin (PDPN), rather than LYVE-1 which may explain the different levels of LYVE-1 expression.To avoid the influence of these variations, we conducted the next analyses based on the actin signal. The observed orientation of the HDLECs was influenced by channel geometry, like for the HUVECs although not completely similar. In the static condition, HDLECs exhibited circumferential orientation in the tubular channel, similar to HUVECs, while they demonstrated a wide distribution of orientations in rectangular channels whereas HUVECs primarily aligned in the main channel direction (Figure 6 c versus Figure 4 a). Under bidirectional flow, HDLECs maintained a predominantly circumferential alignment in tubular channels, whereas they mostly aligned parallel to the main axis of rectangular channels even though the distribution of alignment remains broad (Figure 6 d); this behavior differs from that of HUVECs, which aligned in the direction of the main channel in both channel geometries for bidirectional flow (Figure 6 d versus Figure 4 b). Consequently, the alignment of the HDLECs seems less sensitive to flow and is primarily determined by geometry. Gong et al. applied noncontinuous flow (generated via 2-3 times of media refreshments per day) in a tubular channel and showed that most of the lymphatic ECs were oriented in the flow direction, however still many cells aligned at ∼ 30°-60° or ∼ 100°-120° to the main flow.[30] Even though the wall shear stress values were not reported, it seems that application of wall shear stress resulting from several times media refreshment versus only once a day influences the orientation of the cells. In both static culture and bidirectional flow culture, the number of HDLECs at the top of the channels was lower than at the bottom (Figure 6 e, i). While this difference was minimal in the tubular channel, there was a significantly lower number of cells at the top of the rectangular channel compared to the bottom. Additionally, morphological differences were observed between HDLECs in different conditions (Figure 6 f-h, j-l). The aspect ratio of HDLECs was slightly higher in tubular channels compared to rectangular channels (Figure 6 f, j). Specifically, some cells in tubular channels exhibited elongated shapes with an aspect ratio of around 3 in static (Figure 6 f) and of around 6 in bidirectional flow conditions (Figure 6 j). Furthermore, HDLECs displayed lower circularity in the tubular channels than in the rectangular channels (Figure 6 g, k). While the cell area was lower in the tubular channel than in the rectangular channel under static conditions, there was no significant difference in cell area between the two channel types under bidirectional flow. (Figure 6 h, l).

### 2.5 Endothelial orientation induced by flow conditions influences monocyte rolling

We studied the influence of EC alignment and morphology on the interaction with immune cells, in particular monocytes. To control EC alignment, we cultured ECs on the bottom of a rectangular microfluidic channel, and exposed the endothelium to different flow conditions. Subsequently, monocytes were flown through the channel and their rolling behavior over the endothelial layers with different alignments, was analyzed (Figure 7 a). ECs formed a confluent monolayer inside the channels under all flow conditions, observed from bottom and side views (Figure 7 b and Supplementary Figure 3). Immunostainings of adherent junctions, actin cytoskeleton and nuclei illustrate EC morphological changes due to flow pre-conditioining (Figure 7 b). As shown in earlier results, the static (no-flow) culture condition induced minor expression of actin with limited stress fibers (Figure 7 b-top). Application of continuous unidirectional flow inducing a wall shear stress of *τ*=1 Pa for 24 hours caused ECs to obtain an elongated morphology with aligned actin filaments along the flow direction, and thin distinct cell edges as seen from *β*-catenin expression (Figure 7 b-middle). When pulsating flow was applied, in which the wall shear stress alternated between *τ*=0.5 and 1 Pa at 1 Hz for 24 hours, ECs were again elongated, but cells and actin stress fibers aligned primarily perpendicularly to the flow (Figure 7 b-bottom). This particular cell morphology displayed discontinuous *β*-catenin expression at the cell membranes and upregulated protein expression near the nuclei; these are effects that are also reported in the vascular system.[67, 68, 28]

**Figure 7:**
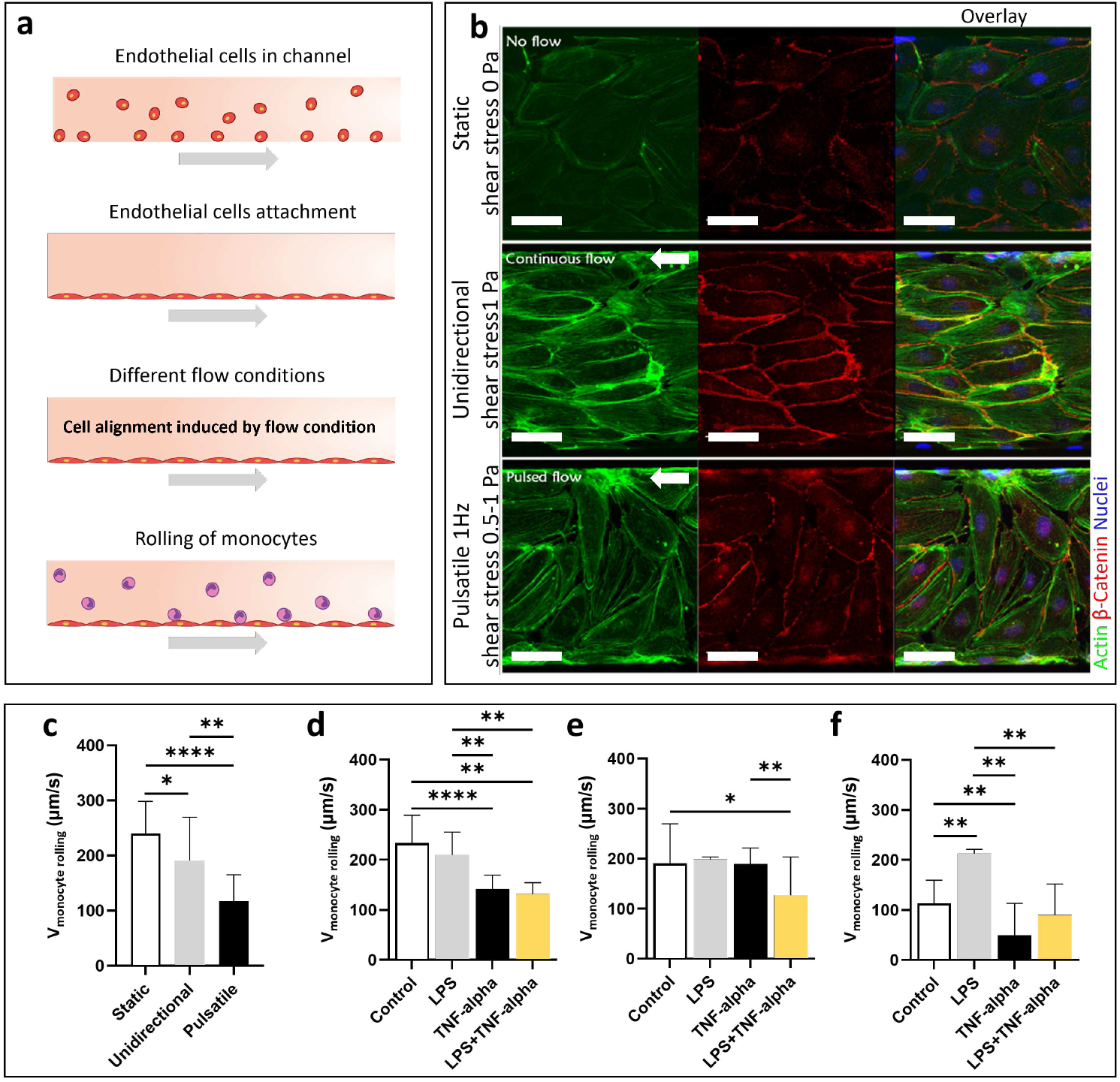
Effect of endothelial alignment on monocyte rolling. a) Schematic representation of the monocyte rolling experiments; ECs were seeded in rectangular microchannels, and the cells attached at the bottom of the channel; different flow conditions were applied to induce different EC alignments; monocytes were flown into the microchannel and monocyte rolling on the endothelial layers was analyzed. b) ECs were cultured under static conditions inducing non-aligned cell morphology, under unidirectional flow at a wall shear stress 1 Pa, inducing cell alignment in the flow direction, and under flow pulsating at 1Hz between wall shear stresses of 0.5 and 1 Pa, inducing alignment primarily perpendicular to the flow. White arrows show the flow direction. Immunofluorescent staining illustrates changes in cell morphology and reorganization of actin and EC junction proteins in response to the different flow conditions. Actin was visualized by Phalloidin staining in green, junction proteins by *β*-catenin staining in red and cell nuclei by Dapi staining in blue. Images from left to right represent actin, *β*-catenin, and the overlay of the two signals. Note: flow was applied from right to left; scale bar: 50 µm. c) Monocyte rolling velocities in the center lane (width: 100 µm) of the channel for the three flow/cell alignment conditions. d-f) Influence of pro-inflammatory stimulation and alignment of ECs on monocyte adhesion behavior; monocyte rolling velocities were determined before (control) and after separate and combined activation of monocytes (by LPS) and ECs (by TNF-*α*) in the three EC alignment conditions of (d) non-aligned cell morphology in static conditions, (e) cell alignment in the flow direction under unidirectional flow at wall shear stress 1 Pa, and (f) cell alignment perpendicular to the flow pulsating at 1Hz with wall shear stress between 0.5 and 1 Pa. *P <0.05, **P <0.01, ***P <0.001, and ****P <0.0001 indicate statistical significance. One-way ANOVA, followed by Games-Howel post-hoc analysis were done to determine statistical significance.

Monocytes in close proximity to the endothelial layer could be visualized rolling over the endothelial layer (Figure 7 a and Supplementary Figure 3). Monocytes were flown over different endothelial morphologies and the rolling velocities were calculated from video recordings (Figure 7 c-f). The parabolic flow profile with corresponding wall shear stress profile inside the wide rectangular channel resulted in faster rolling monocytes in the center and slower rolling cells at the edges of the channel (Supplementary Figure 4); to rule out effects of varying wall shear stress, we considered the monocytes rolling in the middle of the channel only (Figure 7 c). Monocytes rolled fastest on non-aligned ECs and slowest on ECs aligned perpendicularly induced by pulsatile flow (Figure 6 c). The enhanced adhesive behavior on pulsatile flow preconditioned ECs was possibly caused by increased expression of adhesion molecules on these cells. This increased adhesion is in line with the immunostaining observations (Figure 6 b) which suggest this particular cell morphology resembles dysfunctional endothelium, which tends to enhance (pathological) leukocyte recruitment. It is for instance known that ECs from disturbed flow regions have a higher affinity for leukocyte recruitment than ECs from more physiological flow regions.[69] Such healthy microenvironments were best mimicked in our system by the unidirectional flow aligned morphology of ECs[70] and typically show lower leukocyte adhesion at high wall shear stress than at low wall shear stress.[71, 72] Statically cultured non-aligned ECs, on the other hand, did not experience any shear stress prior to monocyte flushing, and consequently shear-stress induced upregulation of adhesion molecules such as ICAM-124 or VCAM-112 was likely to be absent in these cells, possibly explaining the fast rolling of monocytes in this situation.

The influence of chemical activation on monocyte rolling behavior was studied following LPS and/or TNF-*α* treatment (Figure 7 d-f). In the non-aligned endothelial layer, chemical activation of either ECs or monocytes resulted in a decrease in monocyte rolling velocity (Figure 7 d). For this EC morphology, treatment of ECs (with TNF-*α*) had a greater effect on slowing down the rolling velocity than treatment of monocytes (with LPS); treatment of both ECs and monocytes did not cause a further decrease in rolling velocity with respect to the treatment of ECs (Figure 7 d). As discussed above, non-treated monocytes rolled slower on ECs aligned by unidirectional flow than on the non-aligned ECs, indicating enhanced monocyets-EC adhesive interactions on the sheared ECs (Figure 7 c). This effect was, however, slightly recovered after activation of ECs through TNF-*α* stimulation (Figure 7 e). This could indicate an atheroprotective property, which is in agreement with other studies.[73, 74] The combined effect of TNF-*α* and LPS treatment, on the other hand, led to hyper adhesive behavior, which appeared as a sudden drop in rolling velocities of monocytes (Figure 7 e). The same phenomenon was observed on the endothelial layer preconditioned with pulsatile flow; on this perpendicularly aligned cell morphology, where the slowest monocyte rolling velocities were observed, treatment with TNF-*α* alone resulted in a 50% decrease in average monocyte rolling velocity and even monocyte arrest occurred for many monocytes (Figure 7 f). Thus, the effect of TNF-*α* stimulation on monocyte rolling behavior was dependent on underlying EC morphology. On both sheared EC morphologies (induced by unidirectional and pulsatile flow conditions), LPS activation resulted in recovery of reduced monocyte rolling velocity in flow aligned culture conditions (Figure 7 c, e, f).

Activated ECs show an increased expression of adhesion molecules and may also produce chemokines involved in leukocyte trafficking.[75] Although we did not measure the presence of these molecules in our study, their effects may have been stronger in the TNF-*α*-activated ECs that were subjected to shear stress prior to monocyte circulation. In a study of Chiu et al., it was shown that a physiological level of wall shear stress mediates EC responsiveness to TNF-*α* stimulation.[76] While shear stress decreased VCAM-1 and E-selectin expression in response to TNF-*α*, ICAM-1 expression was increased. As ICAM-1 counter LFA-1 is activated in LPS treated monocytes, this combination could substantially influence the adhesion behavior seen in our study. Altogether, our data show that the EC microenvironment plays an important role in the adherence behavior of monocytes to the endothelial layer, which is in line with other studies.[77, 78, 79] Some studies indicate opposite effects with respect to adherence behavior of monocytes to non-aligned versus flow aligned ECs. For instance, Srigunapalan et al. demonstrated increased adhesion of monocytes to no-flow cultured ECs as compared to ECs cultured under shear stress.[80] However, in their study monocyte adhesion to ECs was analyzed during static co-culture, where the emphasis is on stable, permanent, adhesion of monocytes to the EC, most likely involving ICAM-1 and/or VCAM-1. In our study, monocyte adherence behavior was analyzed under flow conditions, where the emphasis is on tethered rolling of monocytes to the endothelium. Here, interaction with selectin may play a more dominant role.

## 3 Conclusion

In this study, we investigated the impact of microchannel geometry and flow conditions on the alignment, morphology and function of endothelial cells within organ-on-chip systems, in particular vessel-on-chip. We fabricated tubular channels with a circular cross-section using sugar 3D-printing, and more commonly used channels with a rectangular cross-section using SLA 3D-printing. Our results highlight significant differences in alignment of the HUVECs (blood vessel cells) depending on the channel geometry. In the tubular channels, the cells demonstrated circumferential alignment, and a continuous circumferential migration along the channel surface in static conditions. This behavior suggests that the curvature and geometrical features of tubular channels influence the cell orientation. In contrast, cells in rectangular channels primarily exhibited orientation in the main channel direction under static conditions. Applying flow in the channels, resulting in wall shear stresses acting on the endothelial cells, induced additional changes in cell alignment and morphology. In general, unidirectional flow induced cell alignment along the main axis of the channel in both tubular and rectangular channels, overruling the circumferential alignment induced by curvature of the tubular channels; pulsatile flow resulted in circumferential alignment in tubular channels, similar to the static condition, and in rectangular channels it mainly induced alignment with the main axis - although the alignment was not as strong as for unidirectional flow. The bidirectional flow condition induced cell alignment along the main axis of the channels for both geometries; however, this effect was more pronounced in the rectangular channel. While the influence of flow conditions was more dominant than the effect of geometry, the impact of geometry remained significant under most flow conditions. Specifically, the proportion of cells aligned circumferentially was consistently higher in the tubular geometry than in the rectangular geometry; especially in pulsatile flow conditions this effect was remarkable. The alignment of the lymphatic endothelial cell type we investigated, HDLECs, also showed dependencies on channel geometry and flow conditions; compared to HUVECs, the HDLEC alignment appeared to be less sensitive to flow and was primarily determined by geometry. Moreover, we observed significant morphological differences (aspect ratio, circularity and cell area) of cells cultured in tubular versus rectangular channels under different flow conditions, for both types of endothelial cells.

We investigated the rolling behavior of monocytes on endothelial layers preconditioned with different flows, including unidirectional and pulsatile flow, which resulted in different EC alignment. In comparison with the no-flow control, the monocyte rolling velocity was reduced on pre-conditioned ECs, especially after preconditioning with pulsatile flow, possibly due to increased expression of adhesion molecules. Furthermore, chemical activation of HUVECs (with TNF-*α*) reduced the rolling velocity of monocytes, while chemical activation of monocytes (with LPS) did not alter the rolling velocity much. Interestingly, this effect was more dominant on ECs preconditioned with pulsatile flow and less pronounced on the ECs preconditioned with unidirectional flow.

In conclusion, this study demonstrates that microchannel geometry and flow conditions both play a crucial role in determining endothelial cell behavior in organ-on-chip systems. These findings provide a strong motivation for the continued development of advanced OoC systems with ever more representative geometrical designs and flow control, to maximize the potential of organ-on-chip to revolutionize biomedical research and personalized medicine by providing more accurate and functional models of human physiology. Future research should explore techniques for integrating more physiological geometrical cues in broader types of organ-on-chip devices to better mimic biological phenomena. In addition, designing for appropriate flow conditions is necessary for enhancing the emulation of *in vivo*-like physiological conditions in healthy and diseased microenvironments of organs. This could be facilitated by integrating more sophisticated and versatile pumping units and sensors on the chip.

## 4 Experimental section

### Cell culture for channel geometry and flow effect experiments

HUVECs (Lonza) and HDLECs (C-12217) were utilized in our experiments. The cell passage procedure was standardized across these cell types, with minor deviations noted where applicable. Cells were expanded on T75 cell culture flasks at 37 ° C and 5% CO_2_, and cultured in relevant culture medium. HUVECs were cultured in endothelial cell growth basal medium 2 (Promocell, C-22211) including supplements (Promocell, C-39211). HDLECs were cultured in endothelial cell growth medium MV2 (Promocell, C-22022) including supplements (Promocell, C-39221) Medium was refreshed every 2-3 days. HUVECs were harvested with 0.05% Trypsin/EDTA and HDLECs were harvested using Accutase (Merck, SCR005)) and centrifuged at 900 rpm (revolutions per minute) for 5 minutes. Cells were suspended at desired density to be used in experiments; both cell types were used in 4 · 10^6^ cells/ml for seeding in microchannels. HUVECS and HDLECs were used between passages four and six.

### Cell culture and chemical stimulation for monocyte rolling experiments

HUVECs culture were cultured as mentioned in the cell culture section, except that HUVECs were cultured in EC-growth medium (EBM-2, LONZA). EC-growth medium was supplemented with Single quots (EGM-2, LONZA). Human promyelocytic leukemia cells (HL-60s) were purchased from the European Collection of Cell Culture (ECACC) and cultured in Monocyte-growth medium (RMPI, GIBCO, Invitrogen) in a humidified incubator set to 5% CO2 and 37° C. monocyte-growth medium was supplemented with 10% (v/v) fetal bovine serum (GIBCO, Invitrogen), 5 mM L-glutamine (GIBCO, Invitrogen), 1% (v/v) penicillin (GIBCO, Invitrogen) and 1% (v/v) streptomycin (GIBCO, Invitrogen). By culturing HL-60s for 4 days in Monocyte-growth medium containing 0.4 mM sodium butyrate (SIGMA-ALDRICH), the cells were differentiated into monocytes. To mimic cell activation relevant for inflammatory conditions, HUVECs and monocytes were stimulated with TNF-*α* and LPS, respectively. HUVECs in the channels were incubated with EC-growth medium with 50 ng/ml TNF-*α* (PeproTech) for 4 hours and then washed three times with growth medium to remove soluble TNF-*α*; monocytes were incubated in monocyte-growth medium with 1µg/ml LPS (SIGMA-ALDRICH) for 15 minutes and then washed three times prior to analyzing the adherence behavior in the microchannel.

### Fabrication of microchannels with circular and rectangular cross-section

Chips with individual channels of circular and rectangular cross-section were fabricated for the EC alignment and morphology experiments. The mold for tubular lumen was obtained using a 3D sugar printer, as explained before.[20] In summary, the sugar fibers were printed with a 250 µm extrusion-based nozzle at 90 ° C.

Fibers were cast in PDMS, until PDMS (Sylgard 184 Silicone Elastomer, Dow Corning, 10:1 elastomer to cross-linker ratio) fully covered them, and the complex was left at room temperature overnight. When PDMS partially polymerized in room temperature, it was cured in 65 ° C at least for 1 hour. The room temperature step was necessary to avoid sugar fiber melting prior to PDMS polymerization. The PDMS layer was immersed in deionized water overnight to dissolve the sugar fibers. A 5 mm PDMS layer was punched with a 4mm biopsy puncher for inlets and outlets, and then was bonded on top of the channel layer within PDMS; For this, the PDMS layers (features facing up) first were both exposed to 20 W air plasma for 30 seconds using a plasma asher (Emitech, K1050X); to achieve full bonding, the assembled complex underwent a thermal treatment at 65 ° C for a minimum duration of 1 hour. The design of the rectangular channel mold was made in Siemens NX (Siemens AG) and then transferred to PreForm software (Formlabs). A durable resin cartridge was inserted into a Low Force Stereolithography 3D (3-dimensional) printer (both from Formlabs), and the printing was run. When the print was complete, the platform was placed in Form-Wash (Formlabs) for 30 minutes, in order to wash the uncured resin in isopropyl alcohol (IPA); the resin then cured in Form-Cure (Formlabs) for 1 hour. The next copies of this mold were fabricated using replica molding as explained before.[81] The resin mold was cast in PDMS, cured at 65 ° C for 1 hour, the polymerized PDMS layer was peeled off, and the inlets and outlets were punched with 4mm biopsy punchers. The channel was closed via bonding to a thin film of PDMS layer. For this, the PDMS layers (features facing up) first were both exposed to 20 W air plasma for 30 seconds using a plasma asher (Emitech, K1050X); to achieve full bonding, the assembled complex underwent a thermal treatment at 65 ° C for a minimum duration of 1 hour. For the monocyte rolling chip, additional to the channel cast, a thin PDMS sheet was made on a smooth surface. Cured PDMS was peeled from the master and fluidic interconnection holes were punched with the use of biopsy needles (diameter 1.2 mm, Harris UNI-CORE). The PDMS devices were irreversibly bonded to the thin PDMS sheet upon treatment with a Corona discharger (Electro-TechnicPrroducts), thus creating the device. To enable side-view imaging of monocyte adhesion to the EC layer, the rough cutting edges of the microfludiic device were repaired by placing PDMS solution on these sides. This created a smooth PDMS wall that did not distort the image. Hence, two observation directions were created: bottom-view and side-view observation. Supplementary figure 3 shows an overview of the production procedure and the observation configurations for monocyte rolling; the channels were 200µm wide, 134µm tall and 2000µm long.

### Chip preparation, loading and flow application for alignment and morphology experiments

The chips were autoclaved for sterility and all the next steps were conducted in a safety cabinet. For static and tilting (bidirectional) culture conditions, the chips were placed in a petri dish (94 mm) for incubation, and a smaller petri dish (35 mm), filled with Phosphate buffered solution (PBS) was placed next to the chip to increase humidity and avoid media dry-out. The channels were coated with 20 µl of fibronectin (50 µg/ml, Merck, F2006-1MG). The coating solutions were aspirated out. Next, 20 µl (4e6/ml) of HUVECs were seeded in the side channels and incubated at 37°C for at least 30 minutes to facilitate attachment to the luminal base. To ensure proper cell adhesion to the side surfaces of the channels, the chips were sequentially rotated 90 degrees to each side for 30-minute intervals; After incubation, culture medium was placed into channels and medium reservoirs to flush out the non-attached cells. Medium was refreshed every day for static and bidirectional culture conditions; media were removed from reservoirs and 100 µl of fresh media were added to the reservoirs on one side of the channels. Media flowed through the channels to the other reservoirs due to the hydrostatic pressure difference caused by the difference in liquid levels in in- and outlet.

The cells were cultured in these chips for 3 days before the application of flow conditions. To generate bidirectional flow, we used the VWR® Signature™ rocking platform shaker, which allows for forward or reverse flow in continuous rocking mode at a maximum angle of 15°The chips placed inside the Petri dishes, were placed on the rocker platform inside the incubator. The maximum rotation angle of 15° and speed of 5 cycles per minute were chosen.

To generate unidirectional flow inside the channels, the ibidi Pump System (ibidi GmbH) was used according to the supplier’s instruction manual. The ibidi black perfusion set was connected to the microfluidic device. From the available ibidi perfusion sets, the black tubing inner diameter was closest to the channel dimensions preventing large pressure gradients within the system. The experimental set-up was connected to the chips and located inside the incubator. The fluidic unit (ibidi GmbH) was calibrated for both types of devices to determine the pressure corresponding to the desired flow rate. Two fluidic units (ibidi GmbH) were required to generate pulsatile flow, one for the continuous unidirectional flow and one to pinch off the tubing at a set frequency. The units were connected to the pump and devices according to the supplier’s instruction manual. A continuous, unidirectional flow was set up first; the fluidic unit (ibidi GmbH) was calibrated for both types of devices based on their calibration curve to determine the pressure corresponding to the desired flow rate. The switching time of the second fluidic unit was set to 0.5 s, corresponding to a frequency of 1 Hz. The flow rate then changes periodically. Calibration was performed with the ibidi black perfusion set filled with HUVECs medium at 37 °C. A timer was used to measure the time it took 2 mL of medium to flow from one syringe, through the device, into the outlet tube. This measurement was performed three times. Based on the calibration curves, the following settings were chosen to achieve a wall shear stress of 0.5 Pa in both tubular and rectangular channels; flow rate of 0.1104 ml/min and pressure of 14.1 mbar for tubular channels and flow rate of 0.0923 ml/min and pressure of 12.2 mbar for rectangular channels. The ibidi fluidic unit (ibidi GmbH) was calibrated for the circular and rectangular channel devices.

### Flow and wall shear stress calculation

The maximum induced wall shear stress on the tubular and rectangular channel wall due to the media refreshment was calculated as following,[82]

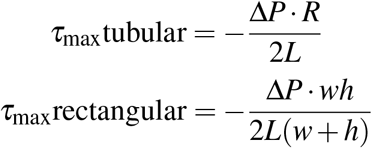

where Δ*P* is for pressure drop between the inlet and outlet of channel at the moment of adding media to the inlet reservoir, *R* is the radius of the tubular channel, *h* and *w* are the height and width of the rectangular channel, and *L* is the length of the channel. We assumed that the flow is fully developed and laminar. Note that the wall shear stress in the tubular channel is spatially uniform and therefore *τ*_max_tubular is the true wall shear stress, but for the rectangular channel the wall shear stress has a spatial distribution and *τ*_max_rectangular is the spatial average of the wall shear stress. The pressure drop generated by media refreshment was calculated by:

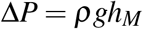

where *ρ* is the density of the cell media (assumed as 1000 kg/m^3^), *g* is the acceleration due to gravity (approximately 9.81 m/s^2^), and *h*_*M*_ is the height difference between the liquid levels at the reservoir and the outlet of the tube. The maximum wall shear stress in the bidirectional flow condition was calculated using the same formulas, however, the tilting rocker movement results in cyclic height and pressure drop variations in time. To consider this, following formulas were used to calculate the time-dependent height change:

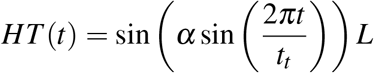

where *α* is the tilting angle of the rocking Platform, HT(t) change in tilting height over time, *t*_*t*_ is the cycle period of the rocking platform, and *t* is time. Hence, the time-dependent pressure difference was calculated using:

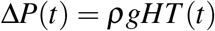

where Δ*P*(*t*) is the time-dependent pressure difference over the channel. The maximum tilting angle of 15° and cycle time of 12 s were chosen. The maximum wall shear stress was calculated at the angle of 15°

The average wall shear stress due to an applied flow rate of *Q* in unidirectional and pulsatile flow conditions for tubular and rectangular channels was calculated by the following equations:

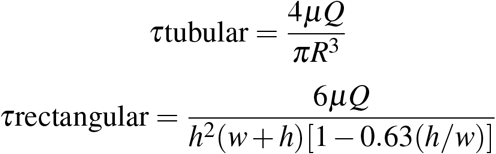

where *µ* is the dynamic viscosity of medium. Again, note that the wall shear stress in the tubular channel is spatially uniform and therefore *τ*tubular is the true wall shear stress, but for the rectangular channel the wall shear stress has a spatial distribution and *τ*rectangular is the spatial average of the wall shear stress.

### Morphology study of ECs cultured in rectangular and tubular channels

The deep learning based image segmentation software Cellpose, developed by Carsen Stringer and Marius Pachitariu, version 2.2.3 was used for HUVECs morphology analysis.[83] With Cellpose we could identify individual cells by using one of the pre-trained models from the Cellpose library. Commonly used models to segment cells are the Cytoplasm (cyto) and Cytoplasm 2.0 (cyto2) models. Both models are trained on two-channel images, where the first is a segmenting channel and the second is an optional nuclear channel.

We used the Cytoplasm 2.0 model because it was also trained with user-submitted images, therefore possibly providing a higher success rate for complex shaped cells. Furthermore, this model can be used as a base model for training with our own images. We used 16-bit RGB color images as input for the Cellpose model. Staining of F-actin (red) provided better contrast in both tubular and rectangular geometries, compared to CD31 (green) for HUVECs and LYVE-1 (green) for HDLECs, and therefore the red channel was chosen as the channel to segment cytoplasm. The blue channel was used for the optional nuclear channel. Cell diameter was set to 70 pixels after calibration with the cyto2 model. The cells in tubular and one rectangular channel images were segmented and where needed, borders or missing cells were manually adjusted. The masks of the segmented cells were saved and the option “Train model with image + mask” with cyto2 model as base model was used. This trained model was then used to identify the cells in all other images. All segmented object outlines were saved as “.zip archive of ROI files for ImageJ”. Open-source software Fiji, version 2.14.0/1.54f was used to show these ROI files and the “Measure” tool was used to generate data on the shape features for individual identified objects. Th orientation polar histograms were generated in Matlab, and morphology violin graphs were generated in Prism-GraphPad.

### Loading of chips for monocyte rolling experiments and morphology conditioning of ECs

The chips were prepared and coated as explained in section ??. The HUVECs were seeded in the microchannels by gently pipetting HUVEC suspension (5e6 cells/ml) into the channels via the fluidic interconnection holes. The cell morphology of HUVECs were influenced by flow. For the non-aligned static morphology, the HUVECs were grown under no-flow conditions. The flow-aligned morphology was achieved by constant flow inducing a continuous maximum wall shear stress of 1 Pa for 24 hours. Pulsed wall shear stress was created by applying a sinusoidal flow, resulting in a sinusoidal maximum wall shear stress ranging between 0.5 and 1 Pa at 1Hz for 24 hours, which caused the perpendicular alignment with respect to flow direction. Flow conditions were obtained by submerging the microfluidic device into a growth medium reservoir and connecting a syringe (BD) with silicone tubing (0.065 inch diameter-standard silicone tubing (HELIXMARK)) to one of the fluidic connection holes. The syringe was placed on a programmable ultra PHD pump (Harvard) that was programmed to withdraw EC-growth medium to flow from the medium reservoir through the channel towards the syringe. Medium reservoirs and chips were kept in a humidified incubator (5% CO2, 37° C) during the flow conditions and the pump with the withdrawing syringe was kept outside the incubator.

### Immunostaining and live imaging

Cells were fixed overnight with 4% formaldehyde for 10 minutes and then permeabilized at room temperature with 0.5% (v/v) Triton X-100 (SIGMA-ALDRICH) in PBS for 10 minutes. Samples were blocked via exposure to blocking solution (1% Bovine Serum Albumin) (BSA) (Sigma-Aldrich, 9048-46-8), and incubated at room temperature for 1 hour. Afterwards the cells were washed twice with washing buffer consisting of 0.05% (v/v) NP-40 (SIGMA-ALDRICH) and 5mM EDTA (SIGMA-ALDRICH) in PBS. *Orientation experiments in tubular versus rectangular channel*. The cell nuclei were stained using NucBlue Fixed Cell ReadyProbe Reagent (Thermo Fisher, R37606) and the cytoskeleton was stained for F-actin using ActinGreen 488 ReadyProbes Reagent (Thermo Fisher, R37110). Staining was performed by exposing the samples to PBS containing 2 drops/mL of each reagent for 60 minutes. HUVECs were stained for CD31 using primary antibody mouse-anti CD31 (Agilent, M0823), and secondary antibody goat anti-Mouse (ThermoFisher Scientific, A28180). HDLECs were stained for LYVE-1 using primary antobody R&D systems (Bio-Techne AF2089) and secondary antibody donkey anti-Goat (ThermoFisher Scientific, A32849). *Monocyte rolling experiments*. To detect adherent junctions, cells were first incubated with primary antibody *β*–catenin (Millipore,1:200 for 1 hour, then washed twice with washing buffer and then incubated with the secondary antibody Alexa555 conjugated donkey anti rabbit (Invitrogen,1:200) for 1 hour. To visualize cytoskeletal actin organization, the cells were incubated with Phalloidin (SIGMA-ALDRICH, 1:200) for 1 hour. Cell nuclei were visualized by staining with Dapi (SIGMA-ALDRICH, 0.1 µg/ml) for 5 minutes. Before and after each of the previous steps, the sample was washed three times in PBS. Fluorescent imaging was performed on either confocal microscope (Zeiss) or fluorescent microscope (Leica-DMi8).The time-lapse videos of the migration patterns of ECs were recorded using inside incubator microscopes (Axion biosystems-Lux).

### Loading monocytes and monitoring the rolling behavior

Monocytes were resuspended in endothelial growth medium and placed into a syringe that was connected to the device via silicone tubing. The setup was placed onto the stage of an inverted microscope within a heating chamber (Zeiss) set to 37°C. The monocytes were flushed over the HUVECs at a rate of 1µl/min and a concentration of 500 cells/µl while monocyte rolling behavior in the plane just above the ECs was analyzed as a measure of initial adhesive interaction with the ECs. The particular low flow speed (corresponding to a wall shear stress of 0.04 Pa in the channel center) was chosen to maximize the amount of visibly rolling monocytes in direct contact with the ECs, while still creating a rolling velocity profile with a steep curve.

A digital HD camera (Sony) was used to record monocyte movement/rolling within a time frame of 10 minutes after starting the flushing of monocytes in the system. The resolution and frame rate were 1440 x 1080 pixels and 25 frame per second (fps) respectively. The observation setup is illustrated in Supplementary figure 3. Single cell rolling velocities and rolling velocity profiles along the width of the channel (x-axis) were determined using video capturing and processing software, VirtualDub (GNU General Public License). Rolling monocytes were defined as cells in focus at the bottom of the channel, which traveled over the monolayer of ECs at a speed higher than 1 µm/sec. The rolling velocity was determined by measuring the rolled distance over several frames and dividing it into time-laps between the first and last frame. The rolling observations were done in a metabolically active system, which include living monocytes, living ECs and warm growth media. Rolling monocytes were analyzed in interaction with the three endothelial morphologies and under four different biochemical stimulation conditions (control, LPS stimulation of monocytes, TNF-*α* stimulation of ECs, and LPS+TNF-*α* at the same time), resulting in 12 experimental groups. Rolling velocities were plotted against the width of the channel (x-axis) to examine the rolling profile.

### Schematics and data analysis

The schematics were made in Adobe Illustrator and Siemens NX (Siemens AG) softwares. Some of schematic items were taken from BioRender library. Imaris software was used for 3D rendering of cell morphology in circular and rectangular cross-section channels.The polar histogram were made in Matlab. The cell tracking was done using TrackMate, an open and extensible platform for single-particle tracking as a plugin in Fiji.[84] Fluorescent images were opened in Fiji and signal brightness was tuned. Cellpose, developed by Carsen Stringer and Marius Pachitariu, version 2.2.3 was used for HUVECs morphology analysis.[83] SPSS software and GraphPad Prism were used for statistical analysis. T-tests and one-way ANOVA tests were done to determine statistical significance; one-way ANOVA, followed by Tukey post-hoc analysis were done to for morphology results, and one-way ANOVA, followed by Games-Howel post-hoc analysis were done for the monocyte rolling experiments. P values <0.05 were considered statistically significant.

---

Supplementary video 1. Time-lapse video of HUVECs at the bottom of channel with circular cross-section in static culture. The track lines show the movement tracks of some random cells in during the time-laps period (22h).

Supplementary video 2. Time-lapse video of HUVECs at the bottom of channel with rectangular cross-section in static culture. The track lines show the movement tracks of some random cells in during the time-laps period (22h).

**Table 1:** Supplementary Table 1. Summary of example studies on endothelial cell alignment, including *in vivo* and *in vitro* (both conventional 2D channels and recent 3D tubular channels). HAECs stands for Human Aortic ECs, HAMEC for Human Adipose Derived Microvascular EC, HPMEC for Human Pulmonary Microvascular EC, HUVEC for Human Umbilical Vein EC, iPSC-ECs for Iduced Ploripotent Stem Cell-ECs, HMVECs for Human Micro Vascular ECs, and HLECs for Human Lymphatic ECs.

| Model (channel type) | Cell type | Flow | Cell orientation | Reference |
| --- | --- | --- | --- | --- |
| *In vivo* porcine iris microvasculature | Porcine eye ECs | Perfused porcine eye | Flow-aligned endothelial cells and concentrically aligned smooth muscle cells | [85] |
| *In vivo* Endothelium in the normal human optic nerve head | Retinal artery and vein ECs |  | Flow-aligned | [86] |
| Well-plate microfluidic devices | HAECs | Unidirectional-flow | Aligned with flow in high wall shear stress | [3] |
| ibidi μslides (Rectangular channel) | HAMEC, HPMEC, HUAEC and HUVEC | Unidirectional-flow | Aligned with flow in high wall shear stress | [4] |
| Cell sheet on thermoresponsive PEG (2D) | HUVECs and iPSC-ECs | Unidirectional-flow | Flow aligned | [87] |
| Vascular network in hydrogel and computational models of cell migration in lumens | hMVECs |  | Circumferential migration in tubular channels larger than capillary size | [40] |
| Lumen in hydrogel-LumeNEXT (bifurcation) | HUVECs |  |  | [14] |
| Lumen in hydrogel | HUVECs, HLECs | Noncontinuous flow by media exchange | Most of the cells were oriented in the flow direction, however, still a lot of cells aligned in the $\sim 30^\circ$-$60^\circ$ or $\sim 100^\circ$-$120^\circ$ angles with respect to flow direction | [30] |
| Lumen in hydrogel | HUVECs | Unidirectional and bidirectional flow | The majority of HUVECs were aligned in $\sim 30^\circ$-$60^\circ$ angle with respect to flow condition.s | [21] |

**Figure 8:**
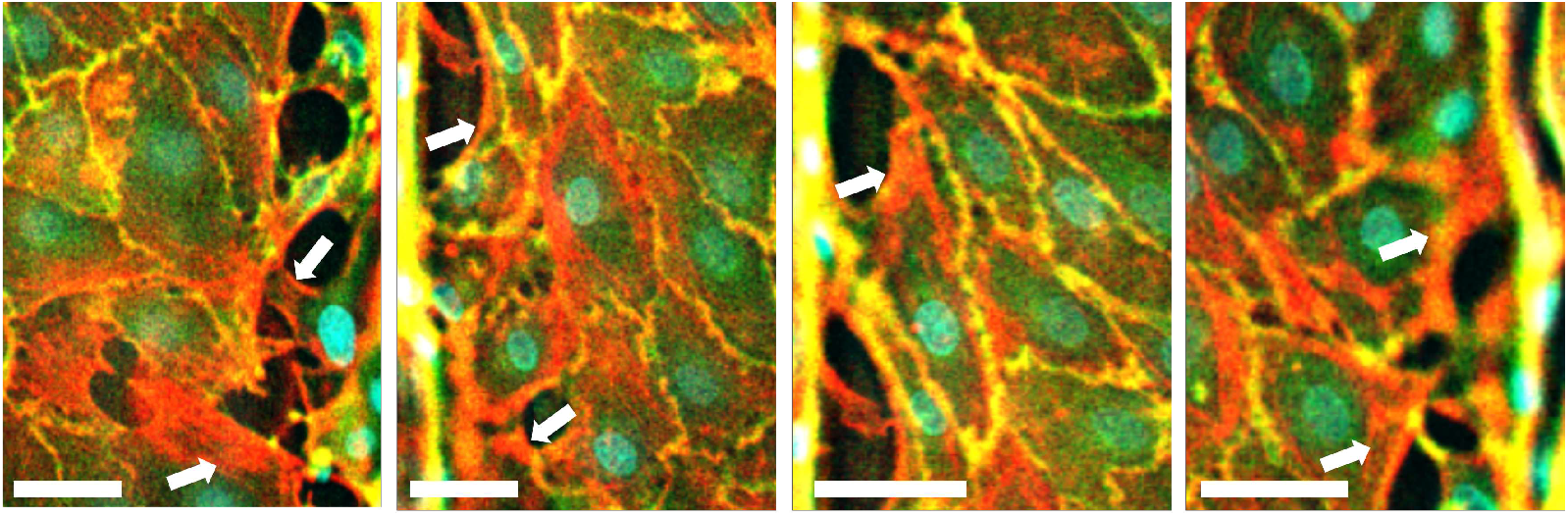
Supplementary Figure 1. Gaps are visible between cells and the channel edges on the bottom of the channel with a rectangular cross-section. Combined fluorescent images of HUVECs stained for cell nuclei (cyan), CD31 (green) and F-actin (red). Prior to staining, the cells had been residing on the bottom the rectangular channel for six days of static culture. Arrows show F-actin stress fibers that appear more intensely in the cells around the gaps. Scale bars, 30 µm.

**Figure 9:**
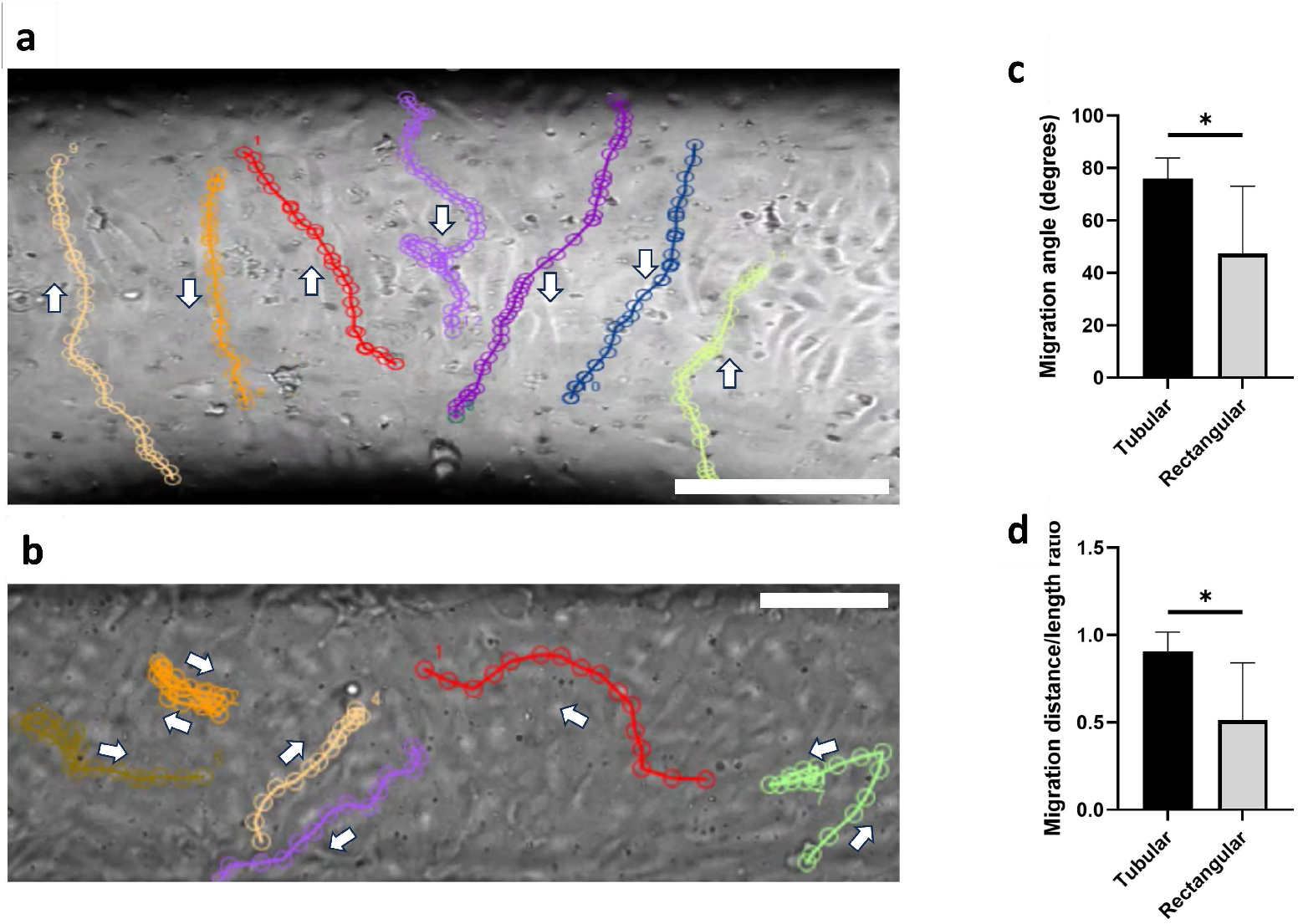
Supplementary Figure 2. Endothelial cell tracking in tubular and rectangular channels. a, b) Phase contrast images of the ECs covering the bottom of the a) tubular and b) rectangular cross-section channels. Shown are the final images of the time-lapse videos shown in Supplementary Videos 1 and 2. The lines in these figures represent the migration trajectory of randomly selected cells in the time-lapse videos; arrows indicate the direction of the motion. Scale bars, 100 µm. c) Migration angle of cells at the bottom of the tubular versus rectangular channels. This angle is defined as the absolute value of angle between the line connecting the starting and final locations of cells after a migration period of 20 h, and the line parallel to the main axis of the channel. d) Migration distance divided by the total length of the trajectory quantifying the straightness of the cell migration, at the bottom of the tubular versus the rectangular channels. For migration along a straight line, this ratio is equal to 1. *P <0.05 indicates statistical significance. Paired t-tests were done to determine statistical significance.

**Figure 10:**
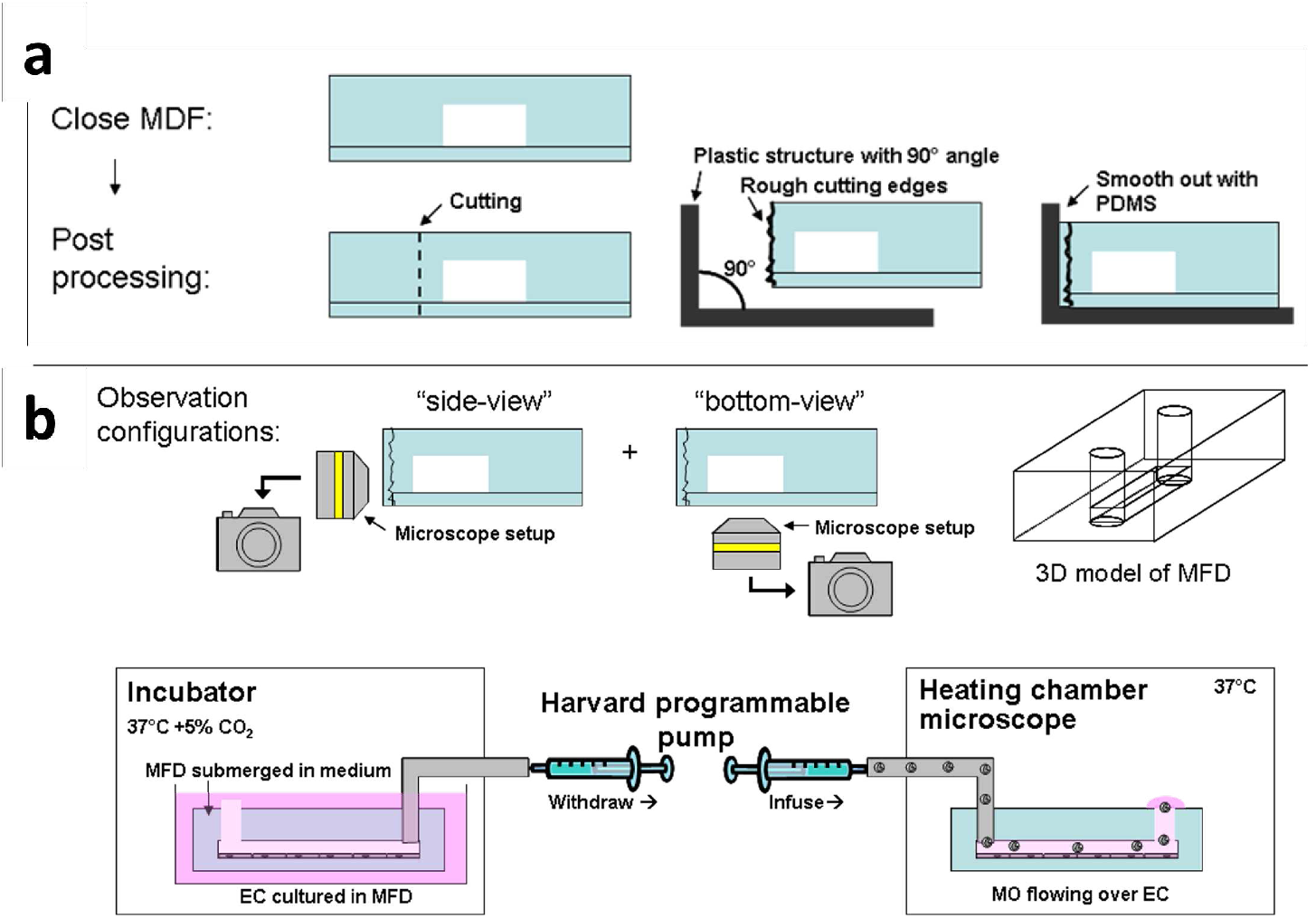
Supplementary Figure 3. Schematic representation of chip preparation and setup for monocyte rolling experiments. a) Post processing required cutting the devices close to the channel and smoothing the rough cutting edges, thus creating a smooth PDMS wall for microscopy to obtain a side-view of chip. This specific processing enabled creating a chip with both “side-view” and “bottom-view” observation possibilities. b) A Harvard pump was used first to create a flow to reach a certain wall shear stress on the ECs cultured in the rectangular channel by withdrawing medium through the channel (left); the chip was located in an incubator to control environmental culture conditions. Secondly, the pump was used to infuse monocytes suspended in medium over the ECs, while placing the chip in a heating chamber to maintain a constant temperature at 37ºC (right).

**Figure 11:**
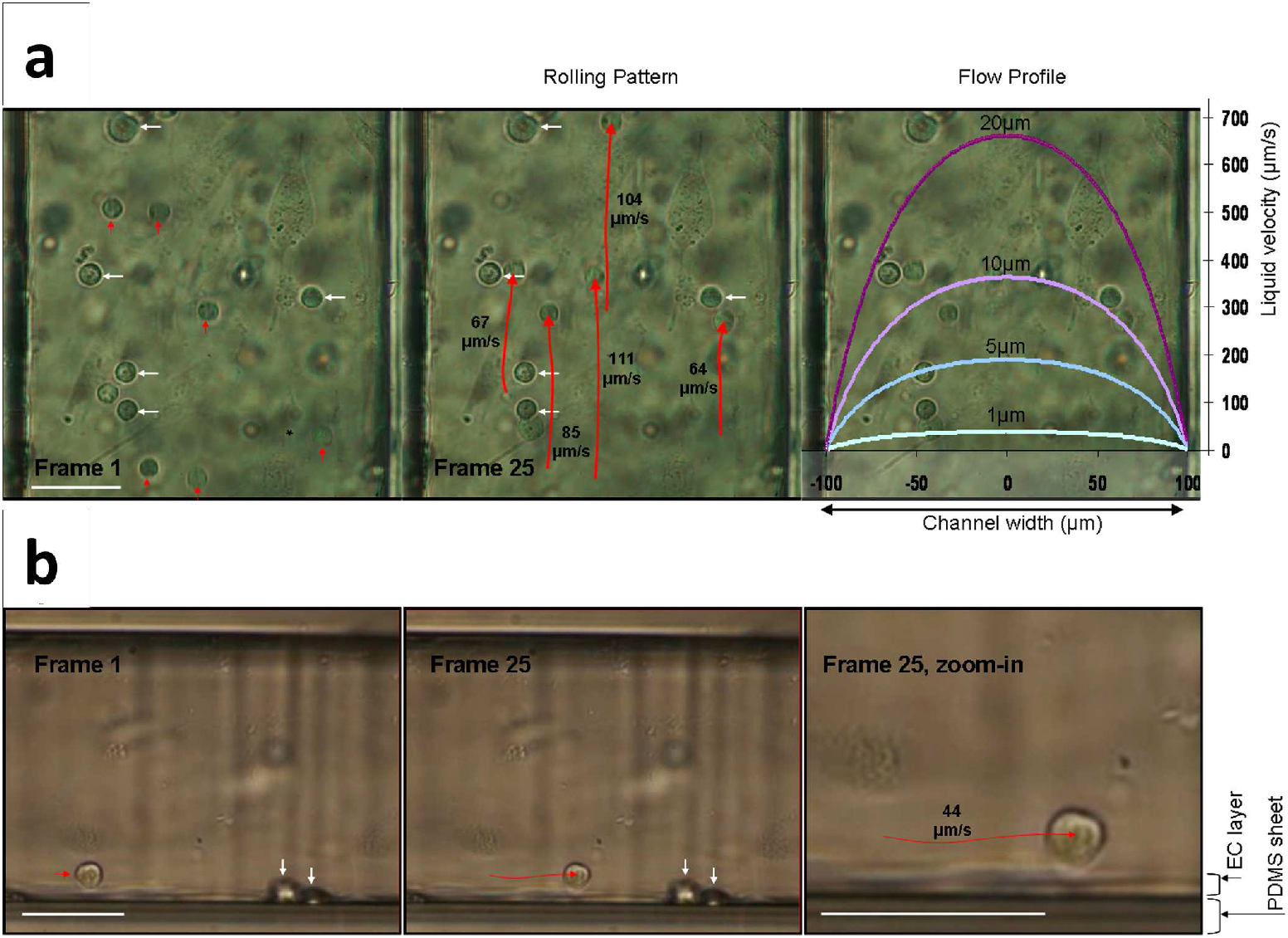
Supplementary Figure 4. Monocyte rolling patterns on ECs are influenced by the flow profile. Inside the microfluidic device, monocytes were flown over a confluent layer of endothelial cells. Video recordings (25 fps) were used to analyze adhesion behavior. Flow direction is from bottom to top in top figures (a, bottom-view), and from left to right in the lower figures (b, side-view). The center of gravity of the cell was used to track movement. In frame 1 and 25, white arrows indicate static adherent monocytes. The red arrows in frame 1 indicate the rolling monocytes and the elongated red arrows in frame 25 indicate the rolling track over ECs. The white scale bar is 50 µm. a) Observation from the bottom-view configuration was used to determine the monocyte rolling velocity. Rolling velocities were higher in the centre of the channel than at the edge of the channel as indicated with the rolling velocities next to the elongated arrows in frame 25. The parabolic flow profile inside the channel which influences the rolling pattern is shown in the rightmost panel. Different colors illustrate the liquid velocities at different heights from the bottom wall. b) From the side-view configuration, monocytes could be observed rolling over the confluent layer of ECs. In frame 25, zoom-in monocyte track was observed going up and down as the monocyte traveled over the cobblestone shape of ECs.

